# Maximum entropy networks show that plant-arbuscular mycorrhizal fungal associations are anti-nested and modular

**DOI:** 10.1101/2025.02.14.637838

**Authors:** Sobia Ajaz, Nida Amin, Álvaro López-García, Henry Birt, Mariona Pajares-Murgó, Luisa Lanfranco, José L. Garrido, Julio M. Alcántara, Matthias C. Rillig, David Johnson, Tancredi Caruso

**Affiliations:** School of Biology and Environmental Science, University College Dublin, Ireland; Department of Soil and Plant Microbiology, Estación Experimental del Zaidín, Spanish National Research Council (CSIC), Spain; Dipartimento di Scienze della Vita e Biologia dei Sistemi University of Turin, Italy; Department of Animal Biology, Plant Biology and Ecology, University of Jaén, Spain; Interuniversity Research Institute for the Earth System in Andalusia, University of Jaen, Spain; Freie Universität Berlin, Institute for Biology, Plant Ecology, Berlin, Germany; Department of Earth and Environmental Sciences, University of Manchester, UK; Lancaster Environment Centre, Lancaster University, Bailrigg, Lancaster, UK

**Keywords:** Plant-AMF association, network structure, null models, maximum entropy bipartite networks, nestedness, modularity

## Abstract

Many applications of network theory to plant-mycorrhizal associations have used a bipartite description, in which one set of nodes is the plants, and the other set is the fungi. Most applications have relied on null models from algorithms that randomly rewire the observed connections to test for non-random patterns in the network. We used existing plant-arbuscular mycorrhizal (AM) fungal datasets to apply a new, well validated generation of network models relaxing the very limiting assumptions of traditional null models. We focused on nestedness and modularity, which have been related to the functioning and stability of communities. Given the existent literature, we expected nestedness and modularity to be prevalent. We modelled plant-AM fungal associations using maximum entropy networks with a degree sequence, soft constraint to generate null distributions for nestedness and modularity. Most plant-AM fungal associations were anti-nested and modular. This pattern was consistent across habitat types and multiple spatial scales. Anti-nestedness can easily emerge from modularity when network patterns are determined by the identity of the plant and AM fungal nodes. Future studies will have to test how the observed patterns determine the ability of the associations to adapt to environmental changes.

## Introduction

High-throughput sequencing techniques have shown that arbuscular mycorrhizal (AM) fungi form intricate, multi-species communities associated with plant species (Öpik et al., 2009; Hart et al., 2015; Morgan and Egerton-Warburton, 2017; Luo et al., 2020). These methods offer detailed insights into AM fungal diversity, detecting not only dominant AM fungal species but also rare or low-abundance taxa that may have been overlooked with traditional approaches. As a result, they highlight how AM fungi can form dynamic and varied associations with plants, influenced by environmental factors (Melo et al., 2019; Lu et al., 2022; Han et al., 2023), geographic location (Ujvári et al., 2021), and the developmental stage of the host plant (Liu et al., 2023).

There is an increasing interest in how AM fungi and plant communities affect each other (Moora and Zobel, 1996; Horn et al., 2017; Wagg and McKenzie-Gopsill, 2023; Ahammed and Hajiboland, 2024; Marrassini et al., 2024) due to the well-established benefits that AM fungi confer on plants, including phosphorus (P) acquisition through hyphal networks (Smith et al., 1997), resistance to diseases (Kaur et al.; Adeyemi et al., 2023; Wahab et al., 2023), defence against herbivory (Babikova et al., 2013a b, 2014), and the broader ecological services of soil carbon accumulation (Hawkins et al., 2023; Wu et al., 2024) and improved agricultural sustainability (Rillig et al., 2019). The latter benefit occurs primarily by reducing the need for chemical fertilizers. One promising avenue of research is the application of network analysis to detect patterns of associations and possible interactions between plants and AM fungi at the community level, which was proposed more than a decade ago but applied only in a few studies (Chagnon et al., 2012; Montesinos-Navarro et al., 2012; Montesinos-Navarro et al., 2019; Sepp et al., 2019; Garrido et al., 2023). These studies have demonstrated the potential of network approaches to reveal complex association structures and their ecological significance, but many questions remain unanswered, underscoring the importance of advancing this approach to better understand community dynamics and ecosystem functioning.

Given the mutualistic nature of most plant-AM fungal associations, a common way to model them is bipartite networks (Figure 1), in which one layer represents AM fungi and the other represents plants, with each node typically representing a species (e.g., Bascompte et al., 2003). However, in cases where AM fungi are identified using sequence analysis, nodes may instead represent virtual taxonomic units (VTUs, defined through molecular data, allowing for a finer resolution of biodiversity rather than species which are taxonomically defined biological units recognized based on interbreeding potential and morphological or genetic traits), allowing for a more nuanced description of fungal diversity (Davison et al., 2015). This approach holds significant promise because it aligns with a robust body of research exploring how the structure of mutualistic networks relates to their stability and resilience (Bascompte & Jordano, 2007).

**Figure 1:**
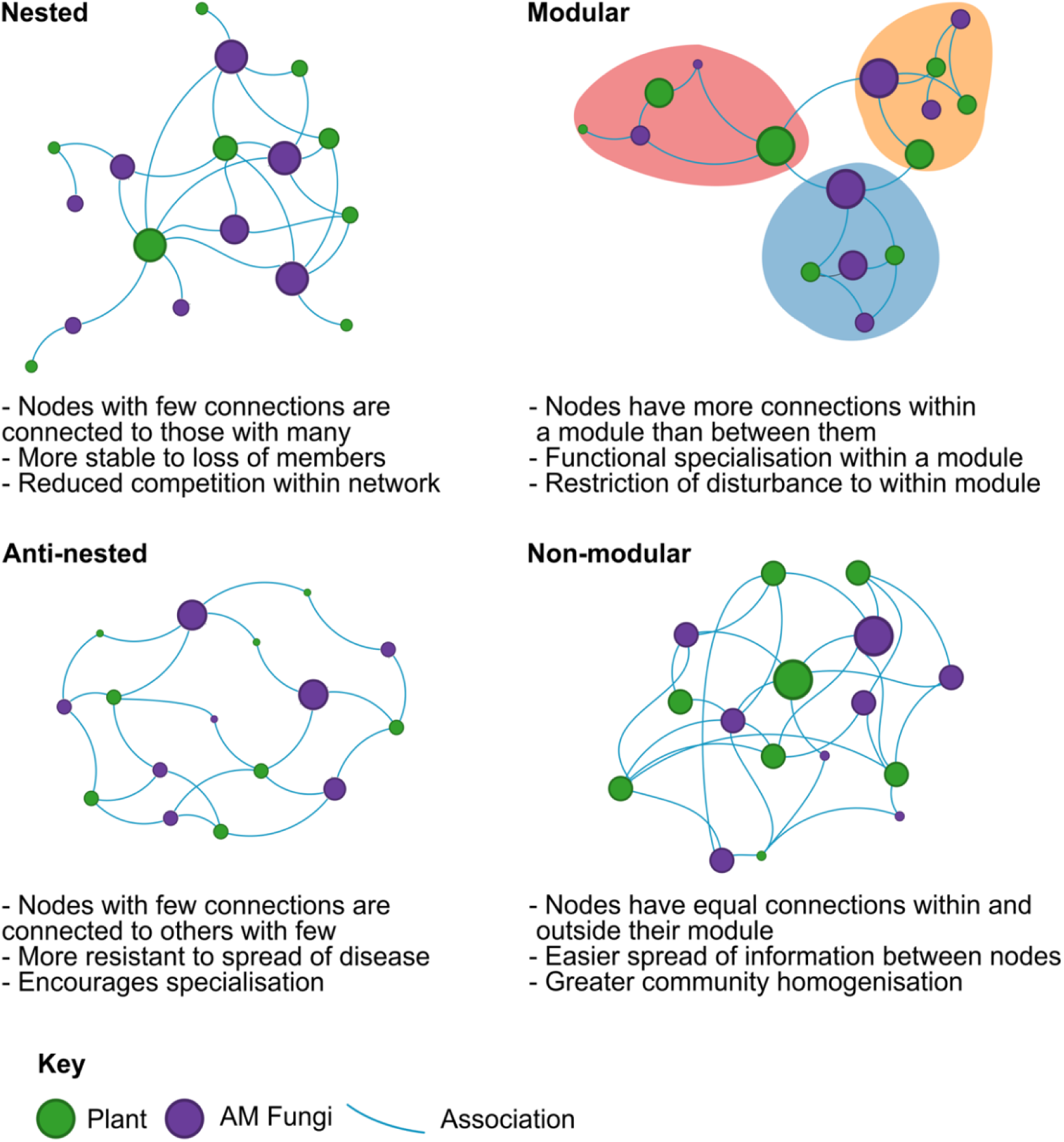
The two possible structures of plant-AM fungal bipartite networks analysed in this work. There are two sets or layers of nodes (Plant species/populations in green, and AM fungal taxa in purple). On the left panels, nestedness and anti-nestedness are illustrated, on the right panel, modularity.

A foundational view in this field suggests that certain network structures are frequently found since they seem to enhance stability, such as by promoting resilience against environmental fluctuations or species loss (Pimm, 1984; Allesina & Tang, 2012). By understanding these structures in plant-AM fungal networks, we can better appreciate the mechanisms that sustain these associations in diverse ecosystems (Stouffer et al., 2005; Montoya et al., 2006). The existing network analyses of plant-AM fungal associations suggest that, despite some complexity due to a relatively high number of nodes and links in some datasets, there are some recurring patterns. Some plants form associations with many AM fungal species, while others connect with only a few. This variability in the number of species connections may lead to the development of nested and modular structures within the network driven by species-specific associations.

Network approaches are thus a useful tool to investigate associations offering potential insights into the ecological and evolutionary dynamics of association patterns (Toju et al., 2014). For example, one possible explanation of nestedness is that the functional similarity (Eisenhauer et al., 2023) of several microbial species relative to their host makes them partially neutral in terms of species identity during the assembly process (Valverde et al., 2020). High nestedness can thus be observed when specialist species interact with a subset of partners with which generalist species also interact, but whether it is related to the stability of mutualistic communities is unclear, with some studies suggesting a positive effect (Bascompte and Jordano, 2007) and others a negative one (Staniczenko et al. 2013).

Another related network structural feature is modularity, describing the tendency for certain species to cluster into communities, or modules, where interactions within each module are more frequent than interactions between modules. This community structure allows for functional specialization within the network, where specific plant and AM fungal species form closer, preferential associations that may have a functional consequence, for example, related to enhanced mutual benefits. In the context of microbial associations, modularity can be contingent on competition and local adaptation (Valverde et al., 2020), which determines the specificity and potential functional complementarity in the degree of association between certain plant species and their microbial symbionts (Guimera and Nunes Amaral, 2005). In other words, a plant community in which each plant species has its unique AM funagl associates, and where thus very few AM fungi are shared between different plant species, would be extremely modular. Since modularity has been observed in plant-AM fungal networks (Montesinos-Navarro et al., 2012; Garrido et al., 2023), it is important to measure its extent and determine the ecological implications of modularity.

The interpretation of the various structural patterns that can be observed in bipartite networks is not always straightforward because, as it often happens in ecology, the same pattern can arise from multiple processes rather than being uniquely associated with one mechanism of formation or a single set of functions. But, at a more fundamental level, the quantification and description of structural patterns in a network are dependent on the application of network modelling techniques. Current techniques mostly rely on null models (Harvey et al., 1983; Gotelli and Ulrich, 2012), which are mostly derived from numerical algorithms that arbitrarily rewire the observed adjacency matrix that codes the information of who is related to whom. The rewiring of the observed matrix is not fully free, as it should be constrained by observed network features relating to ecological and evolutionary factors, such as species-specific interaction preferences, environmental limitations, and functional dependencies. Rewiring must thus satisfy constraints to retain some key aspects of the observed data while randomising all other features related to the identity of the nodes (Bascompte et al., 2003; Blüthgen et al., 2007; Dormann et al., 2009). In other words, the choice of the null model is of fundamental importance for detecting a structural pattern because the model is based on assumptions that affect results, which is important for quantifying features such as nestedness and modularity in ecological networks (Caruso et al., 2022).

We argue that in the case of existing plant-AM fungal datasets amenable to bipartite network analysis, there is a degree of uncertainty in the measurements of plant-AM fungal associations due to a combination of experimental errors inherent to the molecular approaches but also natural fluctuations in populations and associations, both in space and time (Davison et al., 2015; Davison et al., 2018). We thus propose that the best way to assess structural patterns in plant-AM fungal networks is one that models those fluctuations explicitly and we applied that type of model here (see Methods). We identified 35 publicly available plant-AM fungal datasets from different ecosystems, including grasslands, shrublands, forests, and cropland. In those datasets, the AM fungal species are defined either as species or in terms of VTUs (Based on machine-learning models to group sequences or phylogenetic tree) or operational taxonomic units (OTUs, with 97% similarity to species) according to the MaarjAM database (https://maarjam.ut.ee) (Öpik et al., 2010). Specifically, we used a combined maximum entropy and likelihood binary configuration model to analyse plant-AM fungal associations to identify and quantify statistically robust structural patterns of nestedness and modularity that differ from random expectations. We quantified the nestedness and modularity of observed and null model ensemble matrices and tested how universal and widespread nestedness and modularity are in plant-AM fungal association networks. We then offer an interpretation of the functional implications (e.g., ecosystem stability, resilience, and resource efficiency in symbiotic associations) of the observed patterns to identify future research avenues.

## Materials and Methods

### Datasets

the datasets used in the current study were obtained from open-access peer-reviewed publications. We explored all the studies available in the MaarjAM database (https://maarjam.ut.ee) (Öpik et al., 2010) from 2009 to 2019. The most recent publications (2019-2024) were assessed through Google Scholar searches with the keywords ‘plant-AMF association’, ‘AMF abundance” and ‘AMF sequencing in plants”. The 35 different studies were finalised based on the availability of abundance data of AM fungal species in plant roots. AM fungal VTUs/OTUs were treated as species when species-level identification was not possible. The classification was checked through the MaarjAM database (https://maarjam.ut.ee) (Öpik et al., 2010). Some of the datasets were subsets for AM fungi when other fungi were involved in the study, or where AM fungi are found in both plant and soil samples. For this work, the larger datasets were aggregated if multiple levels (treatments, sites etc) of sampling were involved in the dataset (Supplementary Table 1). The datasets chosen possessed a range of AM fungal VTUs (8-277) and plant species (1-245) (Table 1). Data were analysed for at least one interacting plant species where there was a variation across site /replications/ levels/ treatments/ farming systems or different time points.

**Table 1:** Plant-AMF association metadata of the datasets considered in this work.

| *Study* | *No of plants* | *No of AMF species/*<br>*OUT/VTU* | *Sequencing methodology* |
| --- | --- | --- | --- |
| *Alguacil et al., 2011a* | 6 | 12 | Cloning and sequencing |
| *Alguacil et al., 2011b* | 4 | 8 | Cloning and sequencing |
| *Alguacil et al., 2012* | 3 | 20 | Cloning and sequencing |
| *Baar et al., 2011* | 5 | 23 | Cloning and RFLP |
| *Campos et al., 2018* | 3 | 92 | Illumina MiSeq |
| *Davison et al., 2011* | 11 | 40 | Cloning and sequencing |
| *Davison et al., 2015* | 245 | 247 | 454 sequencing |
| *Davison et al., 2018* | 218 | 277 | 454 sequencing |
| *Dieng et al., 2015* | 3 | 13 | Cloning and sequencing |
| *Djotan et al., 2023* | 2 | 321 | Illumina MiSeq |
| *Feng et al., 2021* | 2 | 24 | Cloning and sequencing |
| *Fernández et al., 2022* | 2 | 16 | Illumina MiSeq |
| *Galván et al., 2009* | 1 | 14 | rDNA sequencing |
| *Garrido et al., 2023* | 63 | 87 | Illumina MiSeq |
| *Grilli et al., 2015* | 1 | 59 | 454 sequencing |
| *Guo and Gong, 2014* | 17 | 22 | cloning and RFLP |
| *Higo et al., 2020* | 1 | 148 | Illumina MiSeq |
| *Isobe et al., 2011* | 1 | 24 | Cloning and sequencing |
| *Lara-Pérez et al., 2020* | 1 | 21 | Illumina MiSeq |
| *Li et al., 2014* | 2 | 29 | Cloning and RFLP |
| *Liu et al., 2011* | 2 | 16 | Cloning and RFLP |
| *Moora et al., 2011* | 1 | 71 | 454 sequencing |
| *Moora et al., 2016* | 1 | 47 | 454 sequencing |
| *Muneer et al., 2019* | 3 | 52 | Cloning and sequencing |
| *Öpik et al., 2009* | 10 | 48 | 454 sequencing |
| *Polme et al., 2016* | 5 | 38 | Cloning and sequencing |
| *Sánchez-Castro et al., 2015* | 5 | 37 | Cloning and sequencing |
| *Varela-Cervero et al., 2015* | 5 | 81 | Cloning and sequencing |
| *Sasvári et al., 2011* | 1 | 33 | Cloning and sequencing |
| *Tchabi et al., 2009* | 1 | 49 | Cloning and sequencing |
| *Merckx et al., 2012* | 33 | 56 | Cloning and sequencing |
| *Wang et al., 2018* | 3 | 23 | Cloning and sequencing |
| *Wang et al., 2024* | 12 | 207 | Illumina NovaSeq |
| *Wilde et al., 2009* | 3 | 24 | rDNA sequencing |
| *Zhao et al., 2021* | 10 | 17 | Illumina HiSeq |

The sequencing methods used in the chosen studies also varied. There are 19 studies based on cloning and then sequencing ranging from 2009 to 2018 (including PCR-based amplification of specific regions through RFLP (Restriction Fragment Length Polymorphism)). The other 16 studies are based on Illumina MiSeq/HiSeq/Novaseq, rDNA sequencing and 454 sequencing platforms. The datasets comprise plants from diverse vegetation categories and habitats, including grasslands, cropland, forests, and deserts. As the various studies considered here were conducted at multiple spatial scales, and in some cases with multiple scales within the same study, we consider the scale at which the datasets were aggregated as a key factor in our analysis because aggregation arguably has a major role in both pattern detection and interpretation. We considered three levels: global to very broad scale lists of plant species that may be aggregating plants from different biogeographic regions; more local datasets including the same plant species but under different environmental conditions (either plant populations or experimental treatments); and very local datasets representing physical biological communities as they may be operating on a small scale. This latter case might be considered a local material realisation of associations that are known to be possible given the meta-network of associations in the relevant regional pool of species (Rollin et al., 2024).

To further address in detail the potential effect of data aggregation, we focused on a specific dataset (Garrido et al., 2023, available at Garrido et al., 2024) to validate observed patterns at multiple levels of data aggregation within a single study. This dataset was collected in two different Mediterranean mountain systems in South Spain separated by approx. 100 km. In each system, the plant-AM fungal assemblies were characterised by repeatedly sampling plant roots of several species in different locations of the same ecosystem (between 6 and 24 samples per plant species and mountain system; see (Garrido et al., 2023) for details). Therefore, to further test the impact of data aggregation on the detection of structural patterns and interpretation of those patterns, we considered the following levels of aggregation: plant species meta-network (all locations lumped together), plant species but with the two main mountain systems separately, and population of the same plant species across multiple locations.

### Network modelling

Bipartite networks with two sets of nodes - plant species and AM fungal species - were used to describe plant-AM fungal associations. The abundance (based on the number of reads) of each AM fungal species was converted to a binary matrix (0 or 1 for absence or presence, respectively). The Bipartite R package (Dormann et al., 2009) was used to compute the metrics of nestedness and modularity using the ‘networklevel” function and computeModules function. Nestedness was measured with the nested overlap and decreasing fill (NODF) as per Almeida-Neto et al. (2008). Modularity was measured using the classic approach originally proposed by Newman (2006) but reformulated for the bipartite case by Dormann (2014). The approach uses a binary configuration model to maximise modularity metrics. The maximum value of the metrics is achieved for the partition of the graph into groups of nodes, or modules (i.e. communities) that maximise the number of connections within each module relative to a random configuration model.

To evaluate the influence of both plant and AM fungal species identity on the structural patterns of nestedness and modularity in the association networks, we applied a null model that assumes that the observed association matrix represents the most typical, though not exclusive, network configuration. This comparison helps determine whether specific species identities significantly shape network structure or if the observed patterns could simply result from random associations inherent to some of the statistical properties of the network, as explained below. The network state is defined based on any structural feature relevant to the hypothesis under investigation, and the network is assumed to be fluctuating around an average configuration/state for the measured feature (for example, the total number of links), and any other features dependent on that feature. The structural feature of interest is used as a constraint to build the null model. In our case, given our interest in fully randomising species identity while preserving the number of associations for each species, we followed the classic idea of many ecological null models and used the degree sequence as a constraint.

The degree sequence is the observed number of associations to each species, species by species. But, differently from classic ecological null models, which are based on rewiring algorithms returning random matrices with degree sequences identical to the observed ones, we model the fluctuations around the degree sequence. Following the methodology of maximum entropy models with soft constraints (Squartini and Garlaschelli, 2017), the resulting ensemble of random matrices of our null model respected the degree sequence constraint only on average, which can analytically be achieved by a combined maximum entropy and likelihood approach as synthesised in Caruso et al. (2022) for the case of application to ecological datasets, and documented in details in the references therein (e.g., Cimini et al., 2019). The Python code used to fit the resulting binary configuration model used here is available publicly with examples to run it (Caruso et al., 2022). The full theory behind the model is summarised in Caruso et al. (2022) and references therein.

To compare the observed levels of nestedness and modularity to the null model and make a statement on whether plant-AM fungal associations are nested, modular, or both, we computed nestedness and modularity for each observed matrix. We then fitted the null model to every dataset and calculated nestedness and modularity for the resulting null model matrices. Finally, the Z-score (Neal et al., 2024) of each of the two metrics was computed as (N_observed_ − N_null model_)/SDN_null model_ for nestedness and (M_observed_ − M_null model_)/SDM_null model_ for the modularity metrics. In both cases, 999 sampled random matrices were used to compute the N_null model_ and M_null model_, which, respectively, were the average nestedness and modularity of the null model ensemble. We computed the corresponding standard deviation of those averages (i.e. SDN_null model_ and SDM_null model_).

The Z-score measures the deviation of the observed metric from the null model. For example, if the Z-score of nestedness is negative, the network can be defined as anti-nested, while if positive, it would be nested. In the case of modularity, a negative Z-score would result in a non-modular network, while a positive one would result in a modular network. In other words, our definition of a nested/anti-nested and modular/non-modular network is relative to the baseline of the null distribution. The classic expectation from the literature is nested and modular networks for plant-AM fungal associations, and we thus tested the collection of the analysed dataset for this expectation. Note that, just for simplicity, we define a network as nested/anti-nested and modular/non-modular on the basis of the sign of the Z-score, regardless of p-values or any way to assess ‘statistical significance”. Nevertheless, we also computed the null model p-value for both metrics (Gotelli and Ulrich, 2012; Caruso et al., 2022) to test the null hypothesis that the observed deviation, quantified by the Z-score, is a random fluctuation from the null distribution. A structural pattern can be considered statistically robust or non-random relative to species identity if the observed deviation has a small probability of occurrence (in our case, we chose the widely used p-value of 0.05).

## Results

### Nestedness

Nestedness ranged from 6.33 to 77.30 (Supplementary Table S1), where the minimum value was found for Merckx et al. (2012) and the maximum value was in Zhao et al. (2021). The mean and median values of N_observed_ were lower than the N_null model_ in all except four datasets (Alguacil et al., 2012; Grilli et al., 2015; Higo et al., 2020; Djotan et al., 2023), where N_observed_ was higher. Standard deviation and variances are summarised in Supplementary Table S1. The Nestedness Z-score ranged from 4.75 (nested) to −5.94 (anti-nested), illustrated in Figure 2. A total of 26 out of 35 total datasets deviated from the null model (p-value < 0.05), with 25 datasets anti-nested and 1 (Djotan et al., 2023) nested (see Supplementary Table S1 for a full summary of these results).

**Figure 2:**
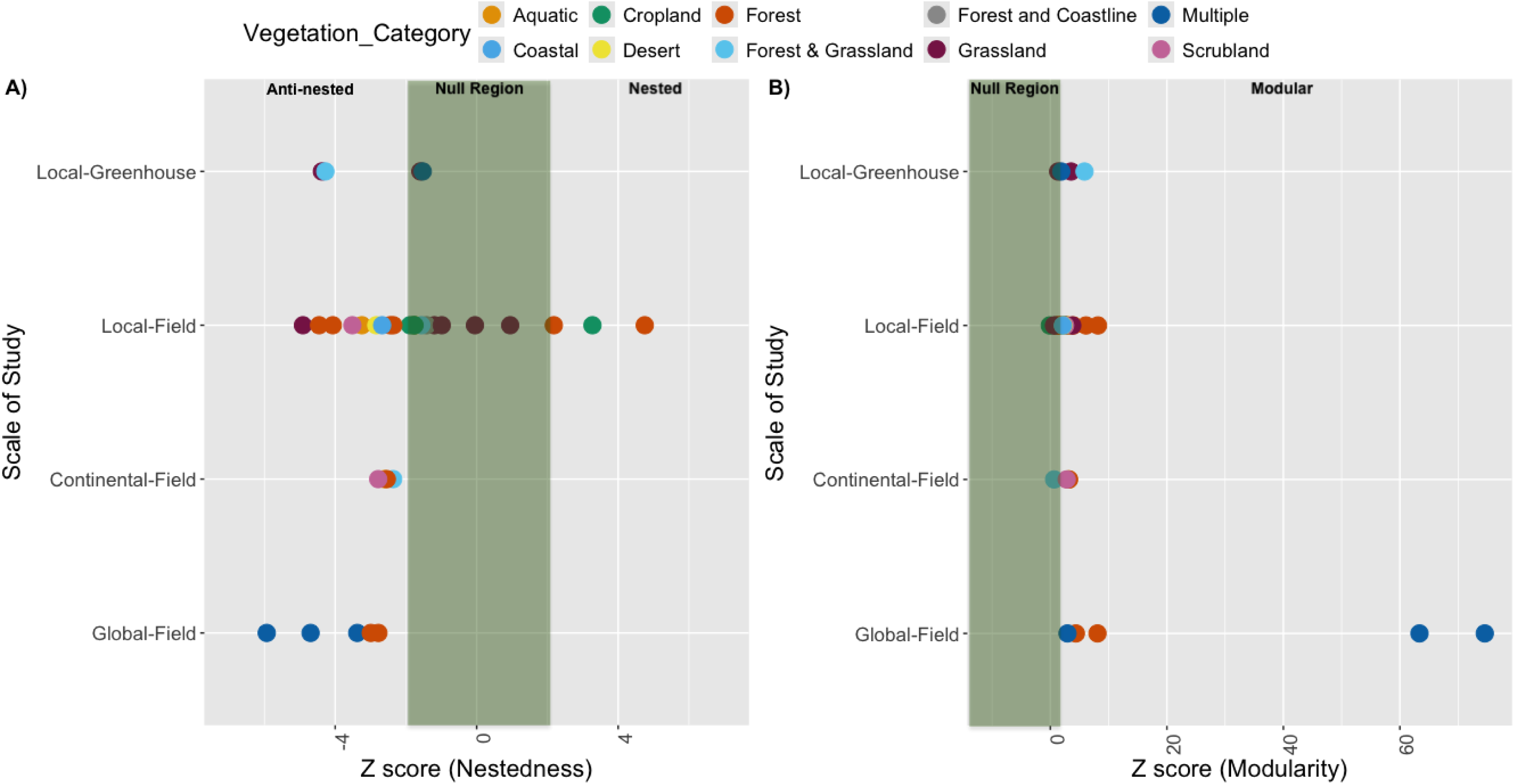
Nestedness (A) and modularity (B) at different scales of study (rows) and across different ecosystems (colours) to profile AM funagl communities in plants. A) Z-score Nestedness is calculated based on N_observed_, average N_null model_ and standard deviation of N_null model._ and B) Z-Score modularity is calculated from M_observed_, average M_null model_ and standard Deviation of M_null model._ The green area represents a Z-score of 0 ± 2 S.E., which under the assumption of a normal distribution of the metric under investigation, would represent approximately 95% of the null distribution: in other words, an observed Z-score falling within the green area is within the 95% confidence interval of the null model expectation for that metric. However, we used this interval in this figure only for visualization purposes: for each dataset, we used the actual null model p-values from the sampled distribution of each dataset specific model fit, without assuming a normal distribution of the metrics. For nestedness (A), a positive Z-score to the right of the green area, that is > 2, implies a nested network (and so <-2 means anti-nestedness, as observed for most datasets). For modularity (B), a positive Z-score > 2 means a modular network, which is what was observed for all networks. The figure thus overall illustrates that almost all datasets were anti-nested and all of them were modular. See Supporting Information for the p-values used in our analysis, and Z-score value distribution.

### Modularity

Modularity ranged from 0.06 to 0.72 (Supplementary Table S1), where minimum value was found in the different tillage plants of maize (Higo et al., 2020) and maximum value was found in herbaceous plant roots (Merckx et al., 2012). Out of the 35 datasets, 26 datasets were found to deviate from the null models (p-value < 0.05), with all of them more modular than the null model (Figure 2). Depending on the datasets, the deviation from the null model could, however, vary considerably. For example, the two largest datasets (Davison et al., 2015; Davison et al., 2018), encompassing a broad range of ecosystems and biogeographical regionals, were found to be exceptionally, highly modular with a Z-score > 63 (p-value <0.001).

### Patterns of nestedness and modularity across vegetation categories

Five datasets with plants distributed across multiple vegetation categories were all found anti-nested (largest Z-scores = −1.53, all p-values << 0.05) and modular (smallest Z-score = 1.88; all p-values < 0.05). The same results were found in the two aquatic plant datasets from lakes (Baar et al., 2011; Moora et al., 2016) showing highly anti-nested (largest Z-score < −2.61, p-value < 0.01) and modular (smallest Z-score > 2.92, p-value < 0.001) patterns of networks with associated AMF communities. The two plant-AM fungal datasets from the coastal areas (Wilde et al., 2009; Guo and Gong, 2014) were found anti-nested and modular as well (largest Z-score NODF <-1.75; p-value <0.03 and smallest Z-score modularity = 2.14; p-value < 0.02). The plant-AM fungal association dataset from the desert (Wang et al., 2018) and the two from Mediterranean scrublands (Varela-Cervero et al., 2015; Polme et al., 2016) were also anti-nested and modular (Z-score NODF < −2.78; and Z-score modularity > 2.53; p-value < 0.01) (see supplementary Table 1 for a full summary of these results).

The eight datasets belong to the forest habitat. The nestedness Z-score values for these forest datasets ranged from −4.46 to 4.75, where six datasets (Öpik et al., 2009; Moora et al., 2011; Merckx et al., 2012; Li et al., 2014; Garrido et al., 2023; Wang et al., 2024) were significantly anti-nested (p-value < 0.01) and 1 dataset (Djotan et al., 2023) was significantly nested (p-value < 0.001). The Modularity Z-score in these forest datasets ranged from 1.29 to 8.17, where six datasets (Moora et al., 2011; Merckx et al., 2012; Grilli et al., 2015; Djotan et al., 2023; Garrido et al., 2023; Wang et al., 2024) were found to be modular and significant (p-value < 0.04).

The cropland vegetation category consists of four datasets, where out of the four cropland datasets, 3 datasets (Galván et al., 2009; Isobe et al., 2011; Sasvári et al., 2011) were found to be anti-nested (largest Z-score NODF > −1.76, p-value < 0.05) and modular (smallest Z-score > 2.19, p-value < 0.02). Only one dataset (Higo et al., 2020) was found to be nested (Z-score = 3.27, p-value = 1) while its modularity did not depart from random expectations.

Plants belonging to grassland were from seven different datasets. Out of the seven grasslands datasets, two of them (Campos et al., 2018; Muneer et al., 2019) were anti-nested (smallest Z-score NODF −4.38; p-value < 0.001) and modular (largest Z-score 3.55; p-value <0.001). The plant-AM fungal interaction datasets from plants’ habitats both in Forest & Grassland show variation in anti-nested pattern and modularity; one out of three datasets (Fernández et al., 2022) was found anti-nested (Z-score = −2.36; p-value <0.01), and modular (Z-score = 6.00; p-value <0.01), while one dataset (Davison et al., 2011) was found anti-nested (Z-score = −2.36; p-value <0.01), and the other (Liu et al., 2011) was found modular (Z-score = 2.40; p-value <0.01).

### Patterns of nestedness and modularity at different levels of aggregation of the same dataset

We initially pooled the whole Garrido et al. (2023) dataset by plant species, regardless of location or habitat differences, effectively constructing a meta-network of all potential associations a plant species might have (Figure 3 and 4). The aggregated dataset was found to be anti-nested (Z-score NODF = −4.10, p-value < 0.05) and modular (Z-score modularity 3.61, p-value < 0.05). The second level of aggregation aimed at identifying local biological communities of plants and AMF effectively associated at a specific geographic location. This was achieved by building one adjacency matrix per mountain system (i.e. Sierra Sur de Jaen and Sierra de Segura). In both mountain systems, Plant-AM fungal association networks were found to be anti-nested (Z-score < −2.18, p-value < 0.01) and modular (Z-score > 2.42, p-value < 0.01) (Figure 3). The third level of aggregation was to account for habitat/dispersal differentiation and local communities associated to different populations of the same plant species and for that we generated adjacency matrices for four individual plant species (*Thymus zygis, Thymus mastichina, Cistus albidus,* and *Crataegus monogyna*) where each location was treated as a different interacting node (Figure 4). All four plant-AM fungal network models were found to be anti-nested (largest Z-score < −2.03, p-value < 0.02) and modular (smallest Z-score > 2.41, p-value < 0.

**Figure 3:**
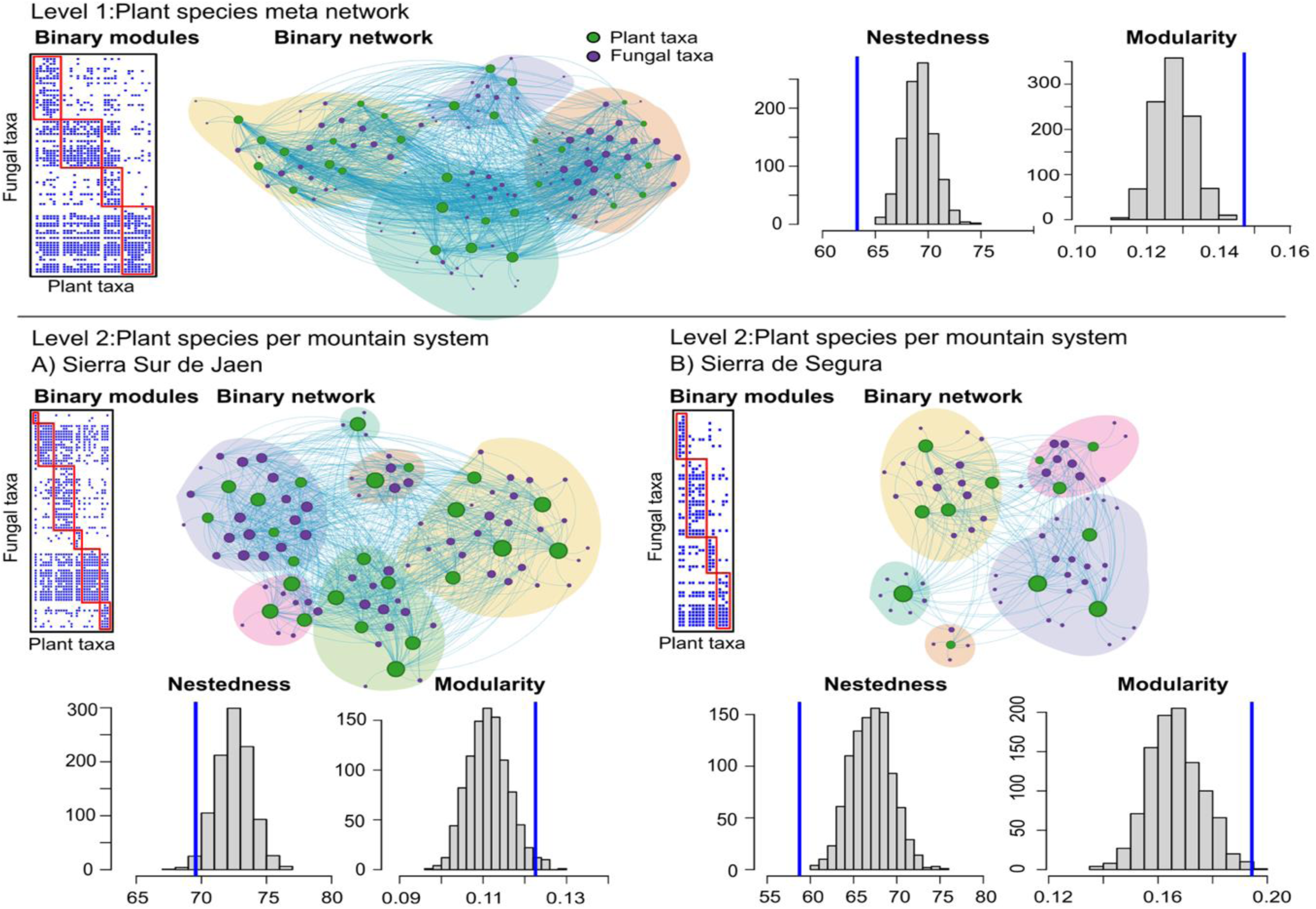
The aggregation of Garrido et al. (2023) datasets at three scales. The first level (top panel, level 1) is plant species, regardless of location or habitat differences. The second level (level 2), is plant species per mountain system with A) Sierra Sur de Jaen and B) Sierra de Segura). See Figure 4 for a further level of aggregation. The figure shows both the matrix and graphic representation of the networks. The matrix is organised by the detected modules (red rectangles), with the blue dots indicating an association between a plant and an AM fungus. In the network graph, the modules are also highlighted by the coloured clouds surrounding the groups of nodes identified in the analysis as modules. The histograms report the results of the null model, with the grey bars representing the frequency (y-axis) of the network metrics (either nestedness or modularity; x-axis) in the null model, and the vertical blue line representing the observed value of the same metric (on the x-axis).

**Figure 4:**
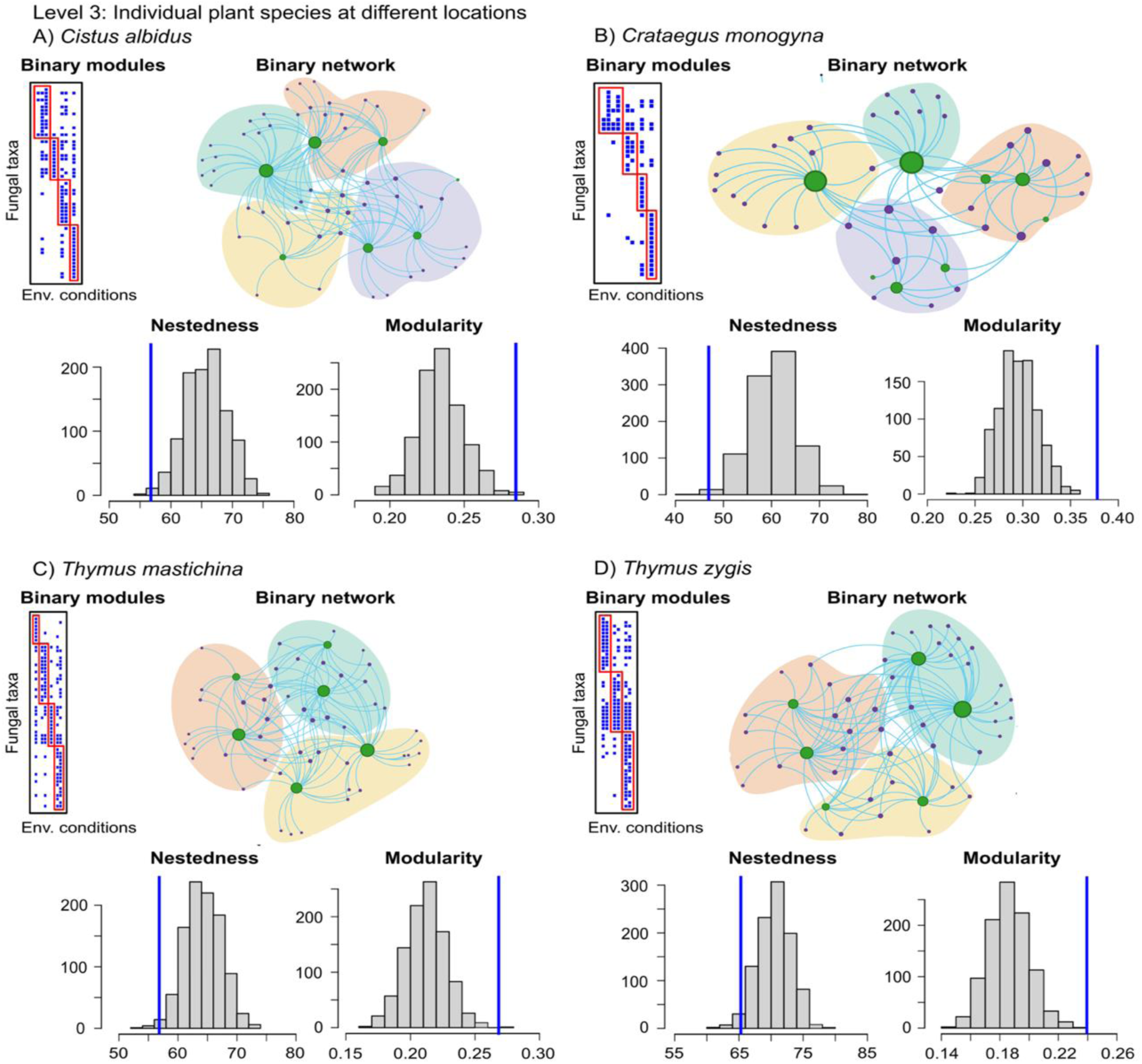
Third level of aggregation of Garrido et al. (2023) datasets (see Figure 3 for the other two levels). Data were here aggregate at the plant species level (*Cistus albidus*, *Crataegus monogyna, Thymus mastichina* and *Thymus zygis*), with the plant node layer representing local population of the species. The figure reports the same type of results as per Figure 3.

## Discussion

Our results show clear evidence of a common set of structural features in plant-AM fungal association networks, namely almost universal anti-nestedness and modularity, which has been found at all the spatial scales and habitat types covered by the studies considered here. As the validation of structural features of networks relies on choices made regarding the null models (e.g. Caruso et al., 2022), we highlight that our modelling approach embraces fluctuations in the major structural features of the association network of plants and AM fungi, notably nestedness, modularity, and degree distribution. These features dynamically respond to environmental and ecological factors, allowing us to assess network features and patterns under varying conditions. This is a new addition to the field of research, and we argue that it is needed for many reasons, which all contribute to the high level of fluctuations in the associations between plants and AM fungi. First, plants and AM fungi interact along a mutualistic-parasitic continuum (Johnson et al., 1997), and along that continuum, individual AM fungal species, populations, and strains can colonise numerous host plants (Sanders, 2002), and similarly, plants may host particular AM fungal species, populations and strains (Kiers and Van der Heijden, 2006). Additionally, certain plant-AM fungal interactions improve plant performance, particularly if the plants are locally adapted (Klironomos, 2003; Koch et al., 2017). Overall, the degree of association between plant species/communities and AM fungal species can vary greatly in space and time at various scales (e.g., see review by (Vályi et al., 2016) and some evidence suggests that stochastic processes may also be a major driver of plant-AM fungal community assembly (Dumbrell et al., 2010; Lekberg et al., 2012).

Fluctuations, which are a combination of natural variability and unavoidable experimental errors such as those inherent to molecular methods, complicate the quantification of patterns that may characterise plant-AM fungal associations and the functional implication of those associations. We therefore argue that our network modelling strategy, in embracing fluctuations in plant-AM fungal associations, offers new insights while also confirming some previous findings. First, and in line with the existing literature, we observed a high level of modularity, which validates the notion that selectivity in plant-AM fungal interactions might result in specific community structures with sets of plant species sharing relatively unique sets of AM fungal taxa (Vandenkoornhuyse et al., 2002; Vandenkoornhuyse et al., 2003; Mony et al., 2021). On the other hand, however, we also observed that plant-AM fungal networks are mostly anti-nested rather than nested, which is contrary to some past observations for mutualistic network. For example, nestedness has been proposed as a common feature of mutualistic networks (Bascompte et al., 2003; Bascompte and Jordano, 2007), including plant-AM fungal networks (Chagnon et al., 2012; Montesinos-Navarro et al., 2012). However, it has also been reported that plant-AM fungal networks can be anti-nested and indeed non-modular (Encinas-Viso et al., 2016), opening the possibility that a high level of nestedness might not be a universal feature of AM associations. Indeed, while we did find some nested networks, we found that anti-nestedness is much more common and coupled with a modular structure in the majority of datasets we analysed. This finding supports the notion that networks with a strong community structure (e.g., highly modular) could be either nested or modular but usually not both (Fortuna et al., 2010). Almeida-Neto et al. (2007) showed that different ecological processes can generate network structures that are less nested (anti-nested) than expected from pure chance. Thus, anti-nestedness should not be interpreted simply as the opposite of nestedness. Moreover, Almeida-Neto et al. (2008) showed that four possible structures (checkerboard, compartmented, beta-diversity, and exclusive subsets) will show lower nestedness than expected by chance. Interestingly, all these structures show a high degree of modularity: checkerboard can be rearranged into a two-module structure, compartmented structures have several modules by definition, while structures defined as beta-diversity, and exclusive subsets have as many modules as plant species; however, in the first case all modules have similar number of partners. Therefore, anti-nestedness and modularity can be interpreted as two ways of quantifying the same type of structure, at least for the case analysed in this work. In other words, if modularity is a key structural feature of plant-AM fungal network, then the same pattern would tend to induce anti-nestedness (see also Figure 1).

How do anti-nestedness and modularity emerge in plant-AM fungal networks and what are the ecological implications? Anti-nested networks can be dynamically stable as shown by some authors (Staniczenko et al., 2013), meaning there could be an eco-evolutionary advantage in the emergence of this structural property in the network. Related to that, and in terms of mechanisms of origin, one hypothesis is that anti-nestedness in plant-AM fungal network could emerge from the balance between mutualism and competition in mutualistic networks (Husband et al., 2002): competition (between plant species as well as between AM fungal species) may shift communities from generalist to specialist association (Ricciardi et al., 2010), while the associations themselves, regardless of how specialised at the species or strain level, may remain mutualistic in terms of the overall function (e.g. exchange of nutrients), which is a benefit to the organisms.

We observed that levels of nestedness and modularity of AM fungal-plant interaction networks can be influenced by several factors. For example, the scale at which the association is investigated—whether in the greenhouse, local, continental, or global field settings—can significantly affect the observed level of modularity, while all networks remain modular. For example, in greenhouse studies (Alguacil et al., 2011a; Dieng et al., 2015; Campos et al., 2018; Fernández et al., 2022) with good control of soil type, AM fungal species, host plants, moisture, and nutrient availability, specific AM fungal-plant combinations can be isolated, which creates specialized modules between treatments in the experiments. In local field experiments, the AM fungal-plant interactions occur in a more natural context under local biotic and abiotic conditions such as in (Galván et al., 2009; Öpik et al., 2009; Tchabi et al., 2009; Wilde et al., 2009; Alguacil et al., 2011a; Isobe et al., 2011; Liu et al., 2011; Sasvári et al., 2011; Alguacil et al., 2012; Guo and Gong, 2014; Li et al., 2014; Grilli et al., 2015; Varela-Cervero et al., 2015; Moora et al., 2016; Wang et al., 2018; Higo et al., 2020; Lara-Pérez et al., 2020; Djotan et al., 2023; Garrido et al., 2023; Wang et al., 2024). In this case, high modularity in plant-AM fungal networks likely arises as a result of environmental heterogeneity and, possibly, also dispersal limitations (Pez et al., 2021; Davison et al., 2015), which would link the network modules to different locations.

We also included continental studies (Baar et al., 2011; Davison et al., 2011; Moora et al., 2011; Polme et al., 2016) that encompass diverse climates, soil types, and geographic features, which increase the assemblage diversity, thereby leading to high modularity resulting from large scale biogeographical and climatic differences in biota. Environmental gradients over large distances are expected to promote unique plant and AM fungal associations, leading to modular structures and, indeed, anti-nestedness because communities with fewer species are not a subset of communities with more species. Over large distances, limited dispersal and local adaptation of plants to native AM fungi can also strengthen modularity, as the association will be more frequent within regional communities than across them. Our study includes also global scale studies (Sánchez-Castro et al., 2012; Davison et al., 2015; Davison et al., 2018; Zhao et al., 2021), which can be explained along the same lines of differences observed at a continental scale.

Overall, the scale of the study is central to interpreting or formulating hypotheses about the observed structures in the network of associations. That was also confirmed by the more detailed analysis of our own dataset published in Garrido et al. (2023), where we could fully control how to aggregate the data at multiple scales from the broad scale of the entire data set to the two different mountain systems included in the study, down to individual plant species at different locations. Modularity and anti-nestedness was observed at all levels, but the interpretation is different: at the highest level, modularity is likely the result of differences due to regional environmental and biogeographical variation. The different regions have unique climatic conditions, soil types, and ecological settings that promote region-specific plant and AM fungal communities, and the modules map onto those regions. At the second level, modularity maps onto two mountain systems, which differ climatically and biogeographically. Additionally, the physical separation of mountain systems restricts AM fungal and plant dispersal, leading to limited cross-system interactions and more localized associations within each system. These factors create distinct modules within the network, reflecting the adaptation of plants and AM fungi to the unique ecological niches in each mountain range. When analysing individual plant species across different locations in a plant-AM fungal network, modularity likely resulted from local populations of the same plant species associating to different AM fungal taxa, which must be due to site-specific factors creating distinct modules.

For very large datasets (Davison et al., 2015; Davison et al., 2018) with > 200 plants, the network showed exceptionally high values of anti-nestedness and modularity, which suggests that clearly defined network structures emerge even in highly diverse sets of species. However, species richness per se cannot entirely explain the observed structure because, otherwise, the null model employed could replicate modularity from the degree sequence constraints, in other words, the observed number of connections to each species, species by species. Our data, instead, fully rejected the null model, meaning that the taxonomic identity of the species plays a significant role in shaping the network (Caruso et al., 2022). While our modelling cannot definitively identify the causes that make plant-AM fungal associations anti-nested and modular, it offers evidence that the key factors are related to the identity of the plant and AM fungal species rather than to the ability of a species to associate with a high or low number of mutualistic partners. Therefore, the processes that structure plant-AM fungal associations must be researched within the eco-evolutionary dynamics that generate diversity between species and their mutualists. These include species-level interactions such as competition within the root, environmental heterogeneity that may allow plant species coexistence at broad scales, and selectivity in plant-AM fungal interactions that can modulate that coexistence as well as plant response to environmental gradients. In other words, taxonomic and population identity matters both for plants and AM fungi and, more specifically, it determines anti-nestedness and high modularity in the investigated networks.

Our modelling of plant-AMF association networks supports strong modularity (always observed) and anti-nestedness (in almost all cases). The robustness of our pattern relies on a network model that relaxes the assumptions of previous null models used to quantify network patterns in plant-AM fungal associations, embracing a major aspect of natural associations: the fluctuations in features such as the number of associations to each node (known as degree sequence). We are therefore confident that our description of network patterns is particularly robust and propose that the next generation of network models applied to any plant-microbial interaction should embrace fluctuations in network properties. Here, we focused on the degree sequence as that is the property considered in the past in ecological null models. Another property to consider in the future is the strength sequence, which is the sum of the weights of all links to a node. Weighted links quantify the strength of associations between two nodes or can also describe fluxes between nodes. Yet, the quest to quantitatively characterise the strength of association between AM fungal taxa and plant species remains an open challenge for the future. Although we cannot offer any definitive conclusion on the reason for which AM fungi and plant association networks have modular and anti-nestedness structure, our cross-scale interpretation suggests that strong network community structure makes plant-AM fungal associations unique and specialised at multiple scales. We speculate that this pattern might allow those associations to better cope with environmental variation. For example, it is known from basic network theory (Squartini and Garlaschelli, 2017; Menczer et al., 2020) that a strong community structure can slow down the speed at which perturbations propagate throughout the network. The application of that principle to ecological networks means that a perturbation of one network module (i.e. a certain plant species and its AM fungi going extinct) will tend to remain confined within that module rather than propagating throughout. A related point is that certain plant-AM fungal associations might be more capable of responding or recovering from certain perturbations and so diversification of the different communities within the network can confer stability to the entirety of the network. Although these ideas are speculative, they are justified by the patterns we quantified in this study but nonetheless require mechanistic, experimental verification in the future.

## Supporting information

Supplementary Analysis S2

## Acknowledgements

The authors acknowledge funding for the European Joint Programme-Soils project ‘Symbiotic Solutions for Healthy Agricultural Landscapes (SOIL-HEAL)’, National support for which came from the German Federal Ministry of Education and Research (031B1266), the Biotechnology and Biological Sciences Research Council (BB/X000729/1), the Department of Agriculture, Food and the Marine (DAFM, project 2021EJPSOILEN303) and Research Ireland (20/FFP-P/8584) in Ireland, and the Italian Ministero delle Politiche Agricole Alimentari e Forestali (project ID170). ALG was funded by MCIN/AEI/10.13039/501100011033 and FSE+ through the grant ref. RYC2022-038499-I (Programa Ramón y Cajal).

## Author contributions

SA and TC conceptualised and designed the manuscript. SA and NA gathered the literature and run the network analysis. SA, ALG, HB, MPM, and TC designed and finalised the figures. ALG, JLG, JMA and MPM perform the aggregation of (Garrido *et al.,* 2023) dataset. SA organized and structured the information and wrote the manuscript with TC. All authors reviewed and contributed to the text of the manuscript.

## Data availability

The data presented in this study are already publicly available as per the references given in Table 1 and Supplementary Table S1

## Supporting Information

Supplementary Analysis 1; Supplementary Table S1.

