## Supplementary Analysis S2 for "Maximum entropy networks show that plant-arbuscular mycorrhizal fungal associations are anti-nested and modular"

### Supplementary Materials:

The following are the datasets used in the network analysis.

The analysis results compiled here include

A) The nestedness is measured with the nested overlap and decreasing fill (NODF). The histogram shows the ensemble values from the null model and the blue bar indicates the observed value.

B) The modularity. The histogram shows the ensemble values from the null model and the blue bar indicates the observed value.

C) The network structure of Plant species

D) The network structure of AMF species

E) Modules formed based on the association between Plant and AMF species

Alguacil, M., Torrecillas, E., Caravaca, F., Fernández, D., Azcón, R. and Roldán, A. 2011a. The application of an organic amendment modifies the arbuscular mycorrhizal fungal communities colonizing native seedlings grown in a heavy-metal-polluted soil. Soil Biology and Biochemistry 43(7), 1498-1508.

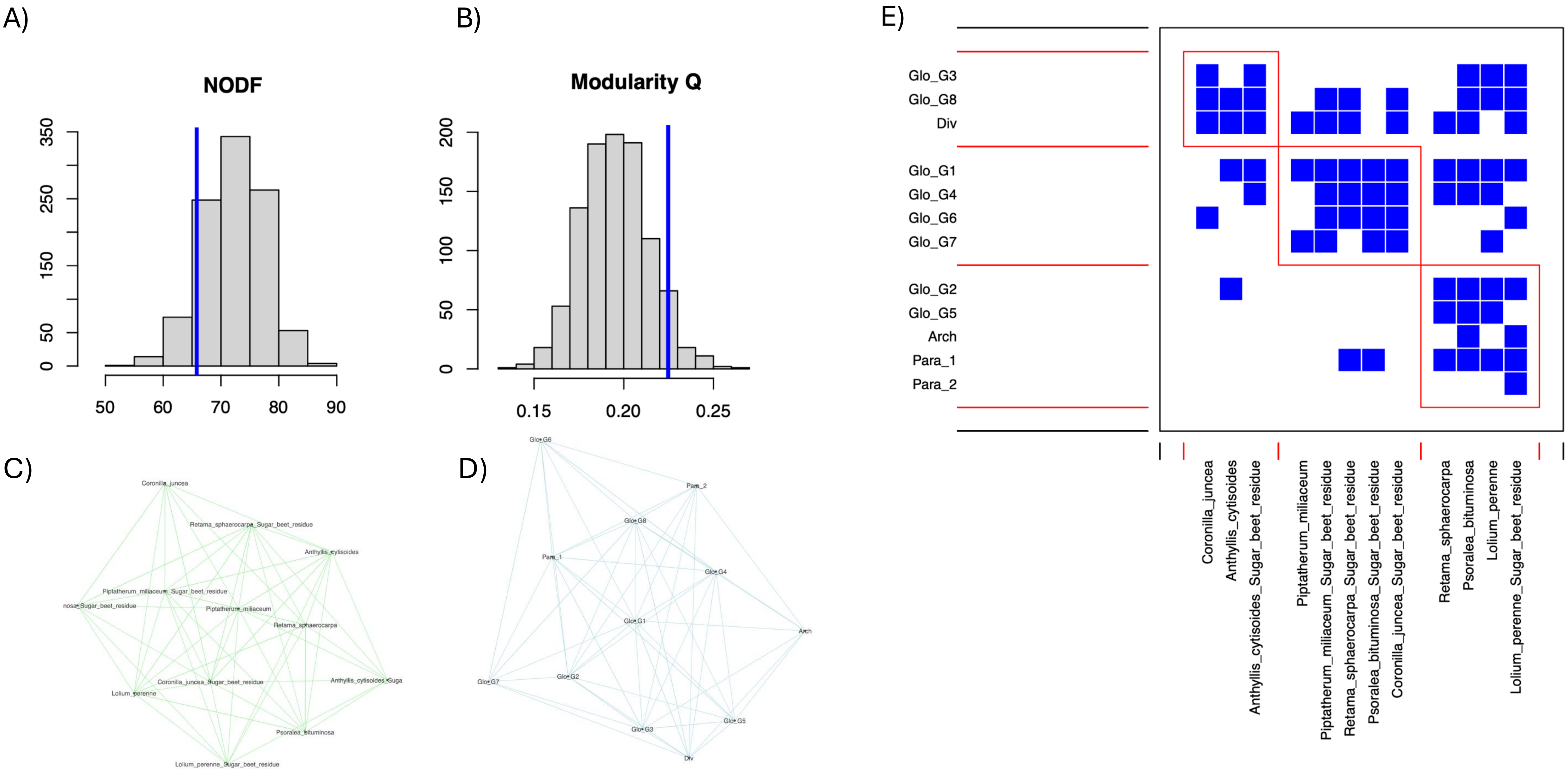

Alguacil, M., Torres, M., Torrecillas, E., Díaz, G. and Roldán, A. 2011b. Plant type differently promote the arbuscular mycorrhizal fungi biodiversity in the rhizosphere after revegetation of a degraded, semiarid land. Soil Biology and Biochemistry 43(1), 167-173.

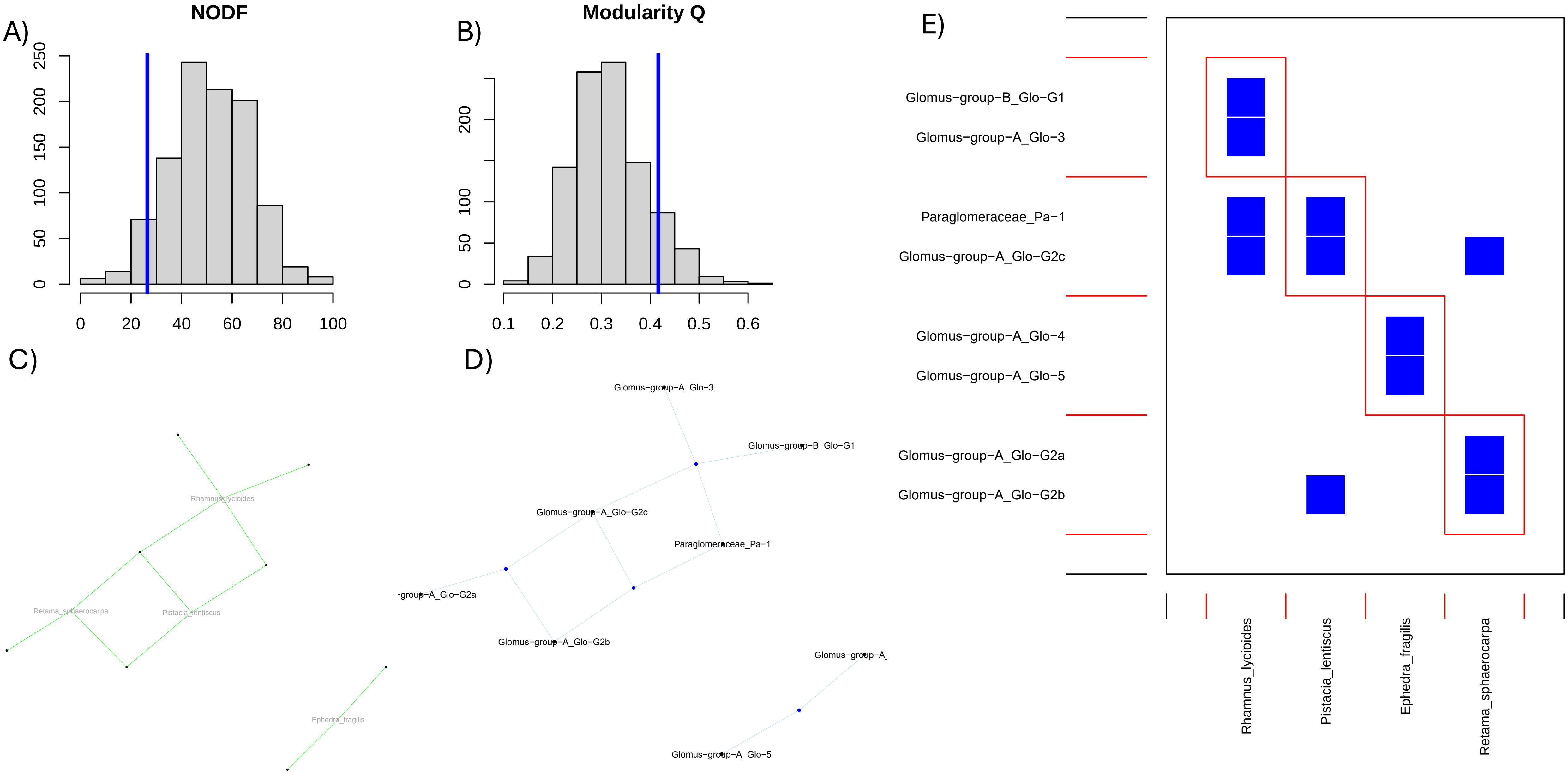

Alguacil, M., Torrecillas, E., Roldán, A., Díaz, G. and Torres, M. 2012. Perennial plant species from semiarid gypsum soils support higher AMF diversity in roots than the annual Bromus rubens. Soil Biology and Biochemistry 49, 132-138.

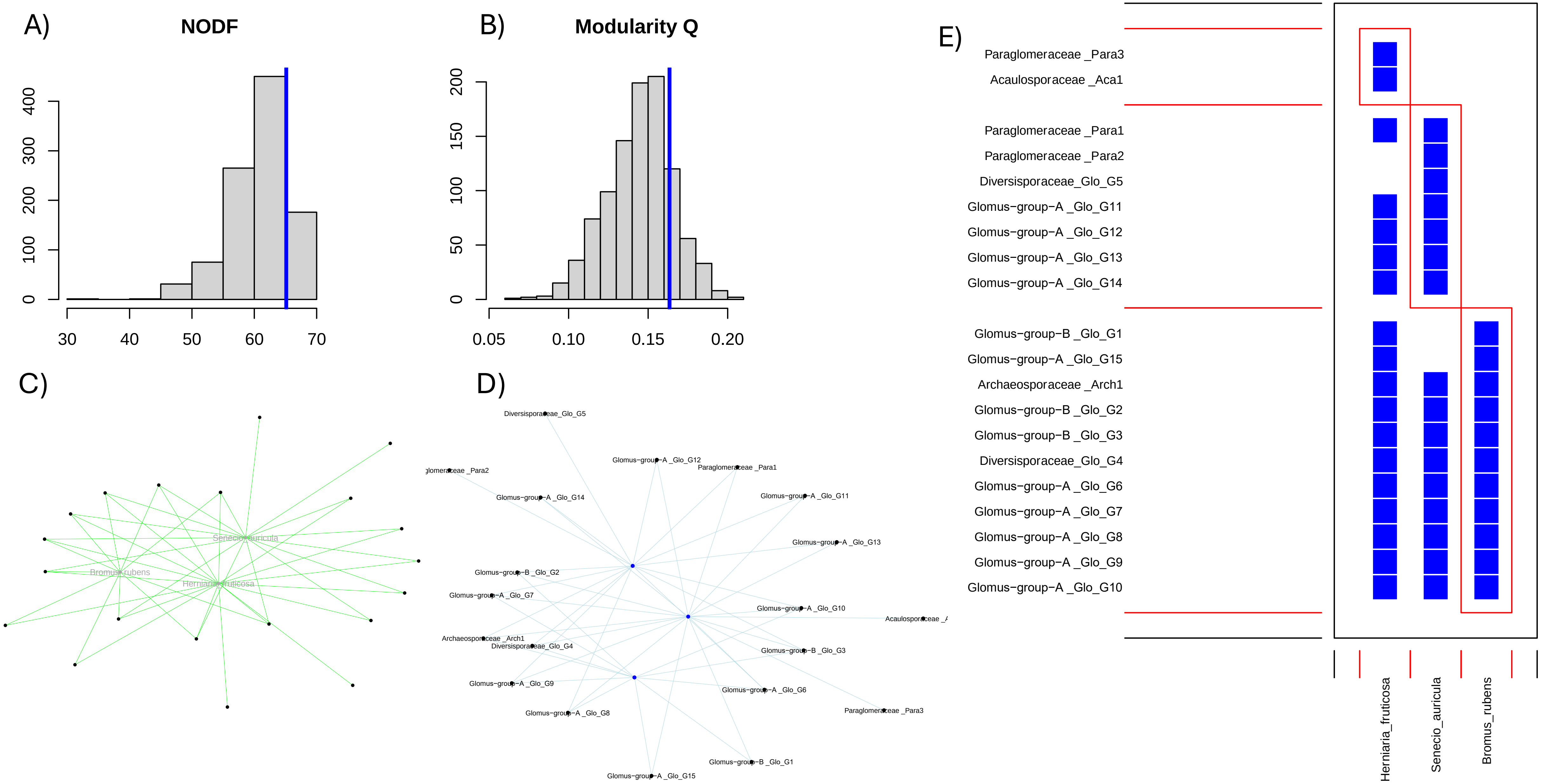

Baar, J., Paradi, I., Lucassen, E.C., Hudson-Edwards, K.A., Redecker, D., Roelofs, J.G. and Smolders, A.J. 2011. Molecular analysis of AMF diversity in aquatic macrophytes: a comparison of oligotrophic and utra-oligotrophic lakes. Aquatic Botany 94(2), 53-61.

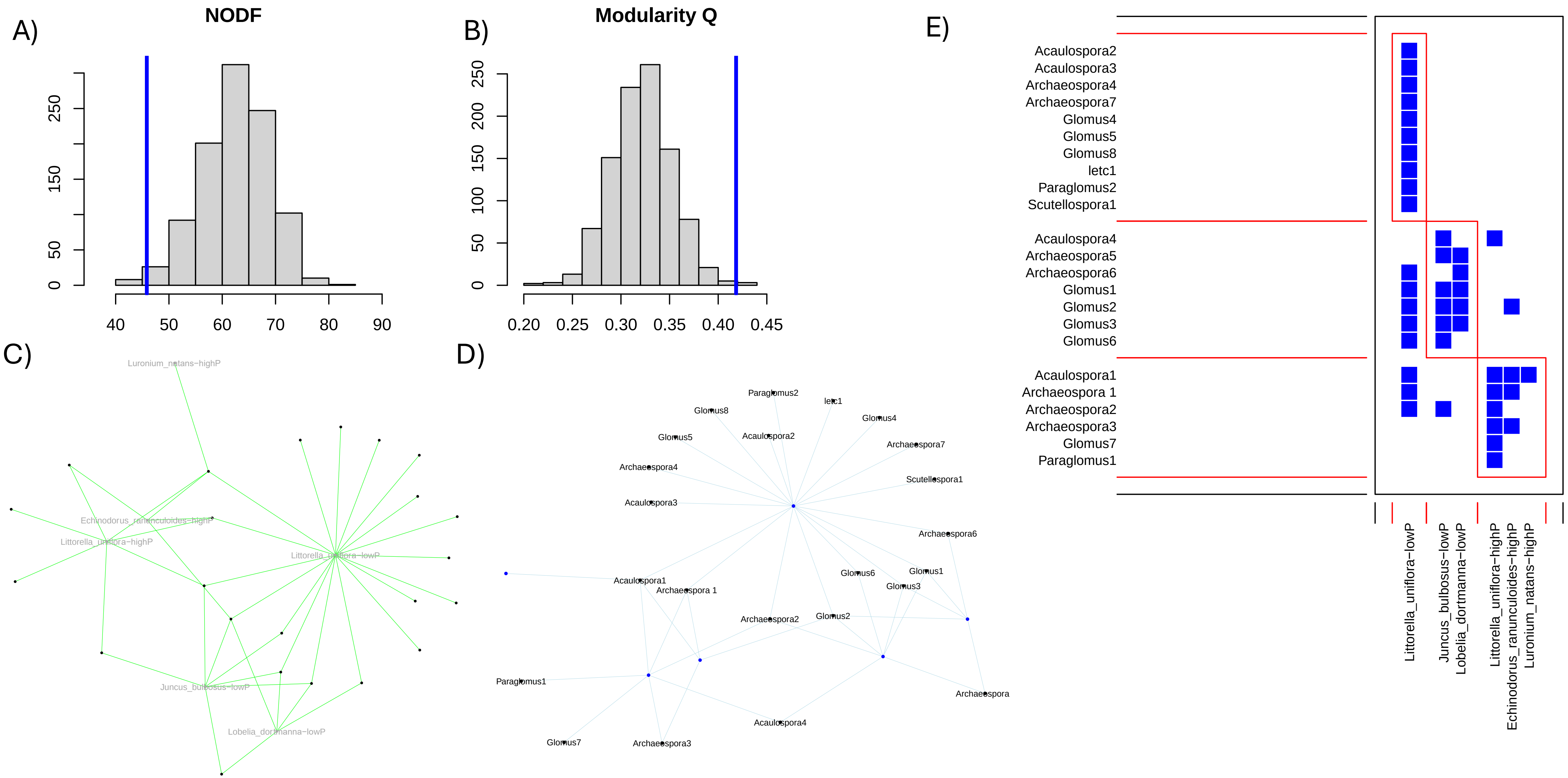

Campos, C., Carvalho, M., Brígido, C., Goss, M.J. and Nobre, T. 2018. Symbiosis specificity of the preceding host plant can dominate but not obliterate the association between wheat and its arbuscular mycorrhizal fungal partners. Frontiers in Microbiology 9, 418365.

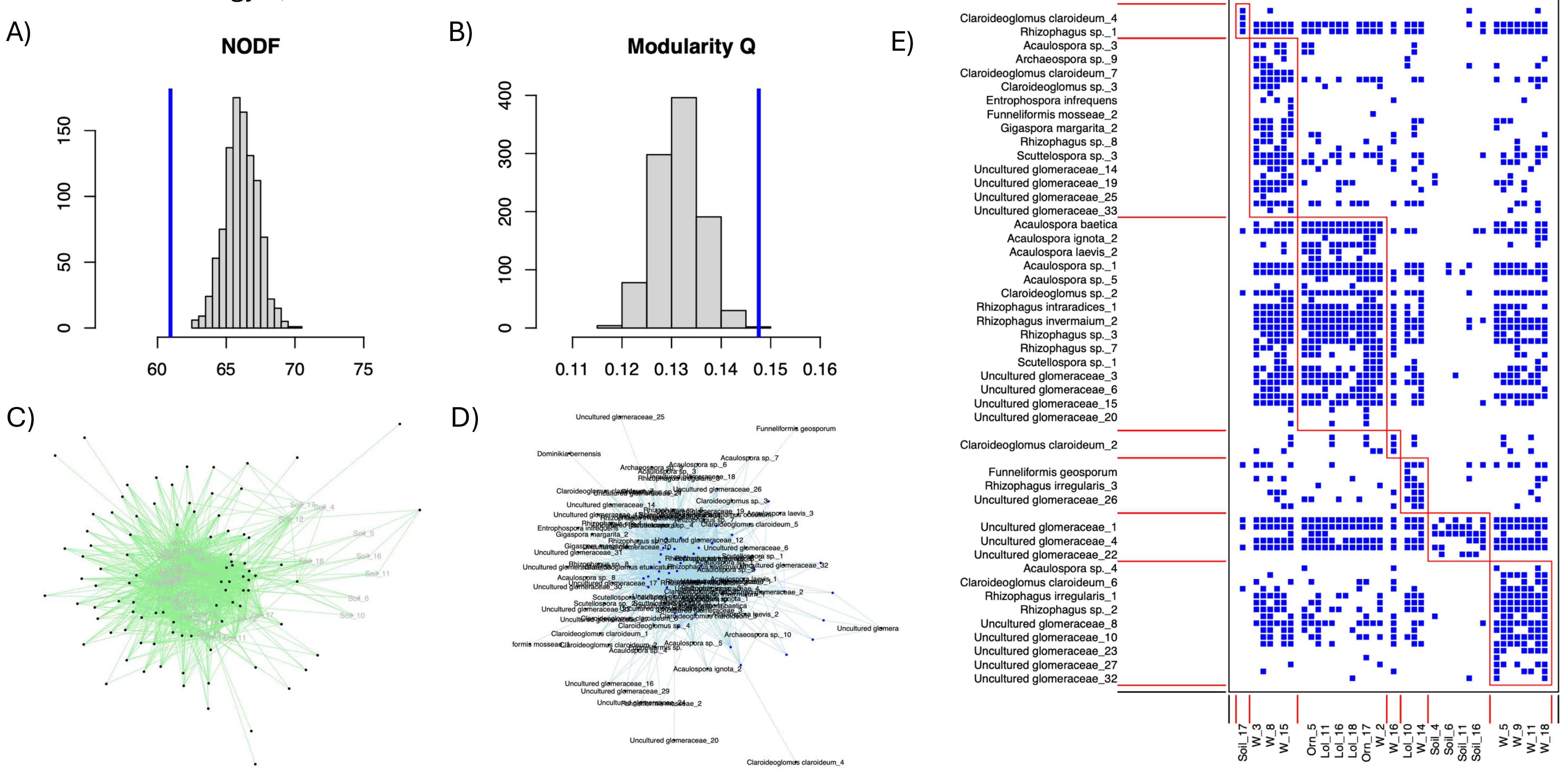

Davison, J., Öpik, M., Daniell, T.J., Moora, M. and Zobel, M. 2011. Arbuscular mycorrhizal fungal communities in plant roots are not random assemblages. FEMS Microbiology Ecology 78(1), 103-115.

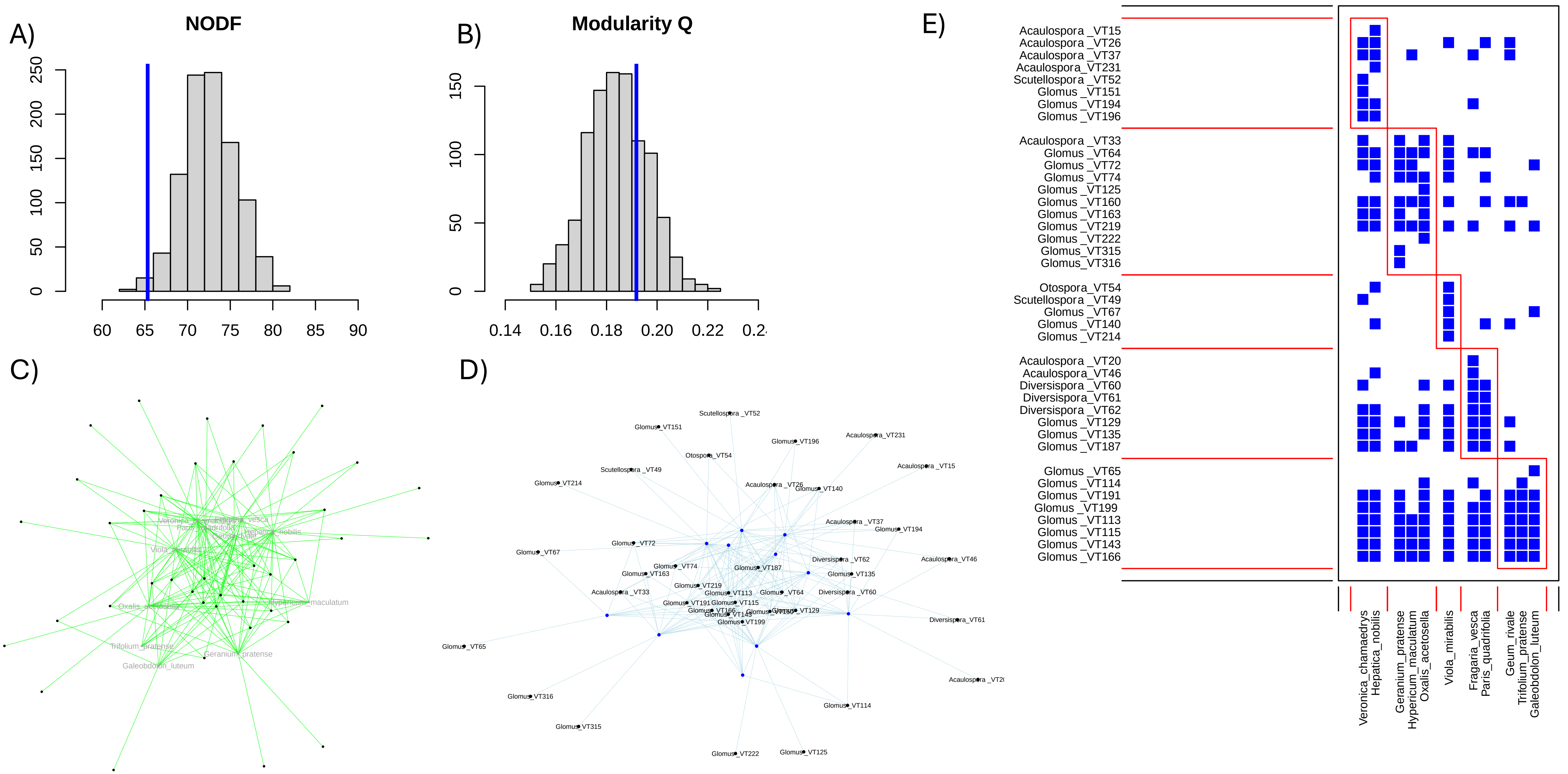

Davison, J., Moora, M., Öpik, M., Adholeya, A., Ainsaar, L., Bâ, A., Burla, S., Diedhiou, A., Hiiesalu, I. and Jairus, T. 2015. Global assessment of arbuscular mycorrhizal fungus diversity reveals very low endemism. Science 349(6251), 970-973.

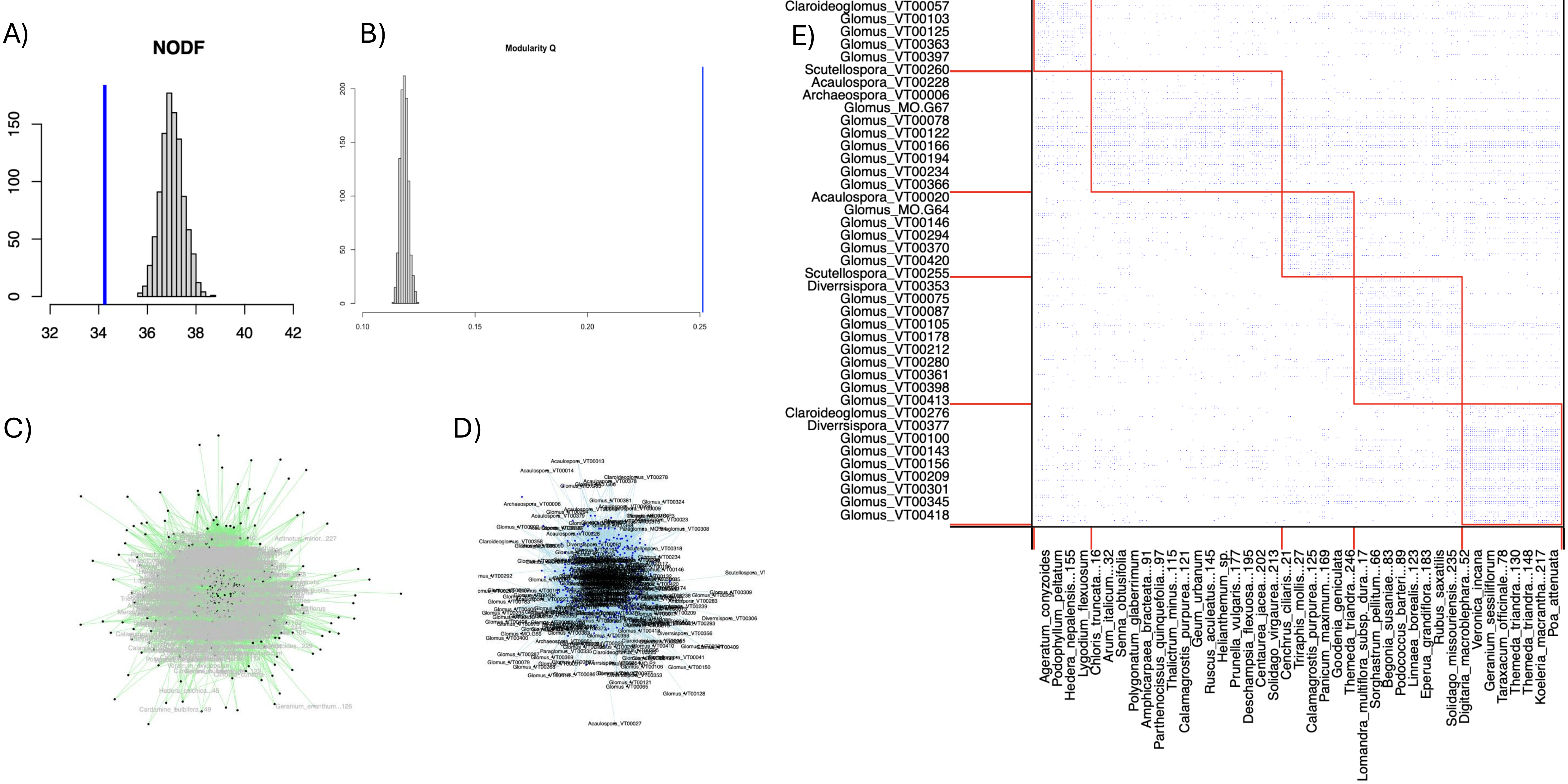

Davison, J., Moora, M., Öpik, M., Ainsaar, L., Ducousso, M., Hiiesalu, I., Jairus, T., Johnson, N., Jourand, P. and Kalamees, R. 2018. Microbial island biogeography: isolation shapes the life history characteristics but not diversity of root-symbiotic fungal communities. The ISME Journal 12(9), 2211-2224.

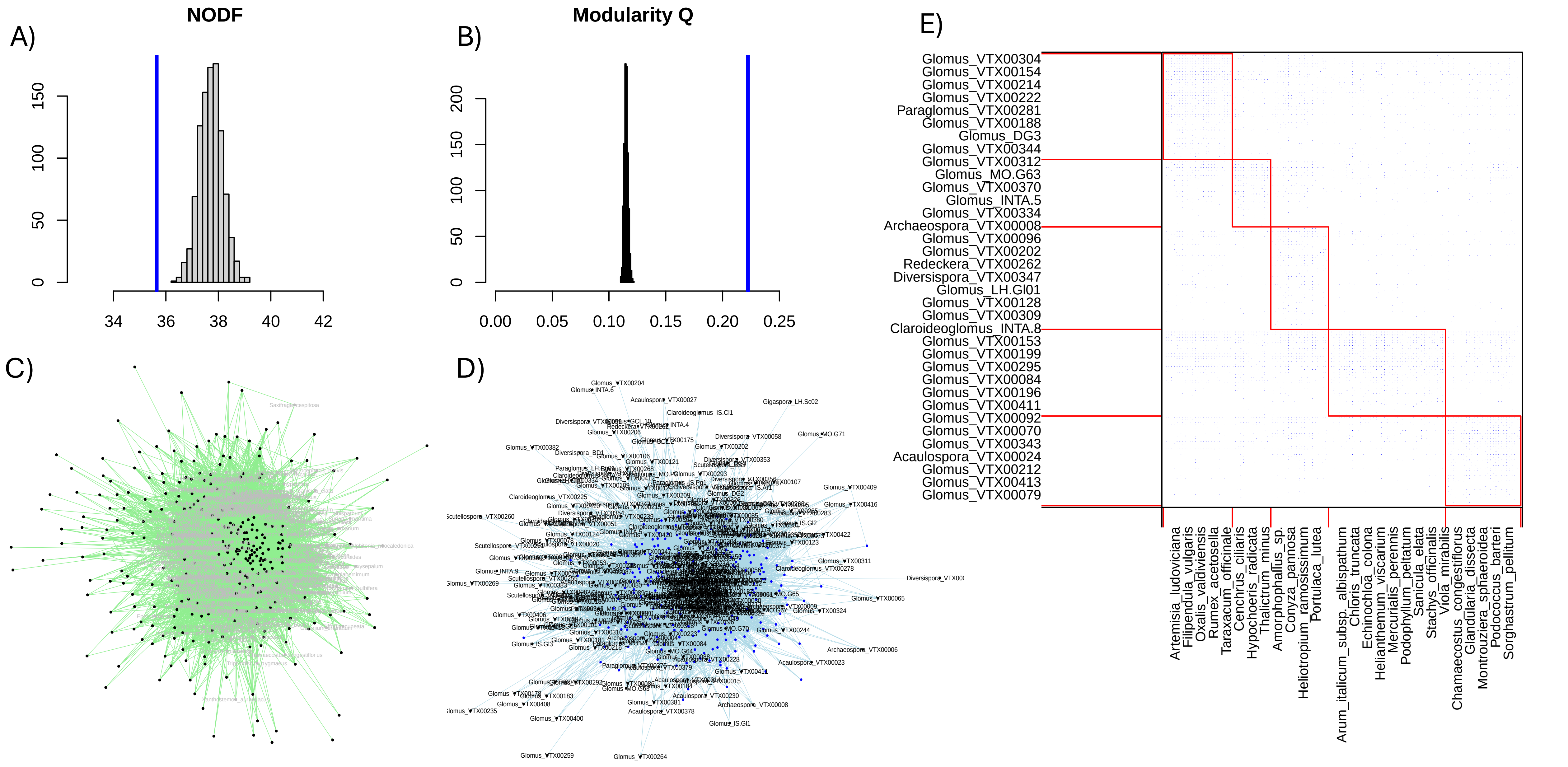

Dieng, A., Duponnois, R., Ndoye, I. and Baudoin, E. 2015. Cultivation of *Jatropha curcas* L. leads to pronounced mycorrhizal community differences. Soil Biology & Biochemistry 89, 1e11.

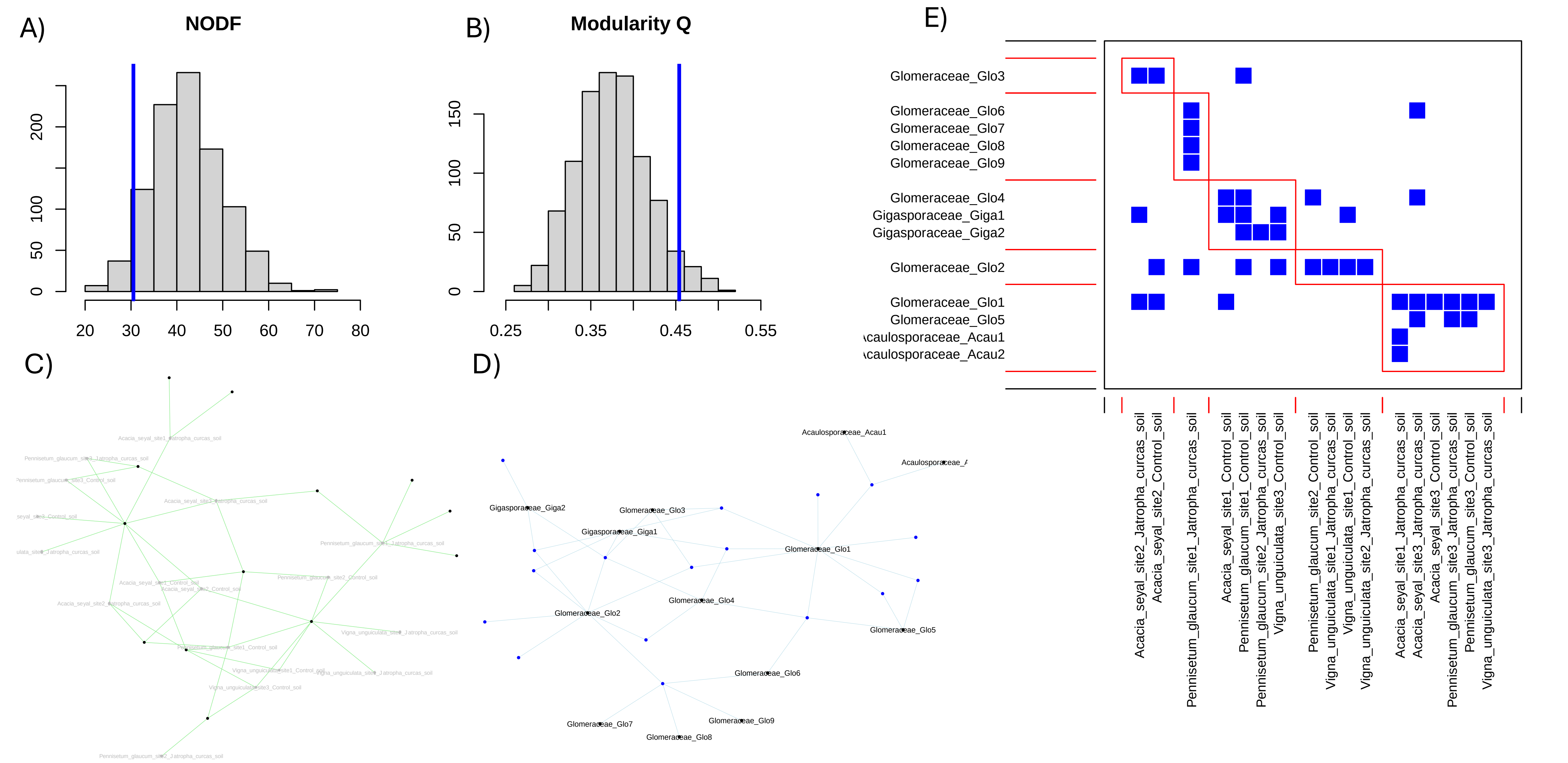

Djotan, A.K.G., Matsushita, N. and Fukuda, K. 2023. Paired root-soil samples and metabarcoding reveal taxon-based colonization strategies in arbuscular mycorrhizal fungi communities in Japanese cedar and cypress stands. Microbial Ecology 86(3), 2133-2146.

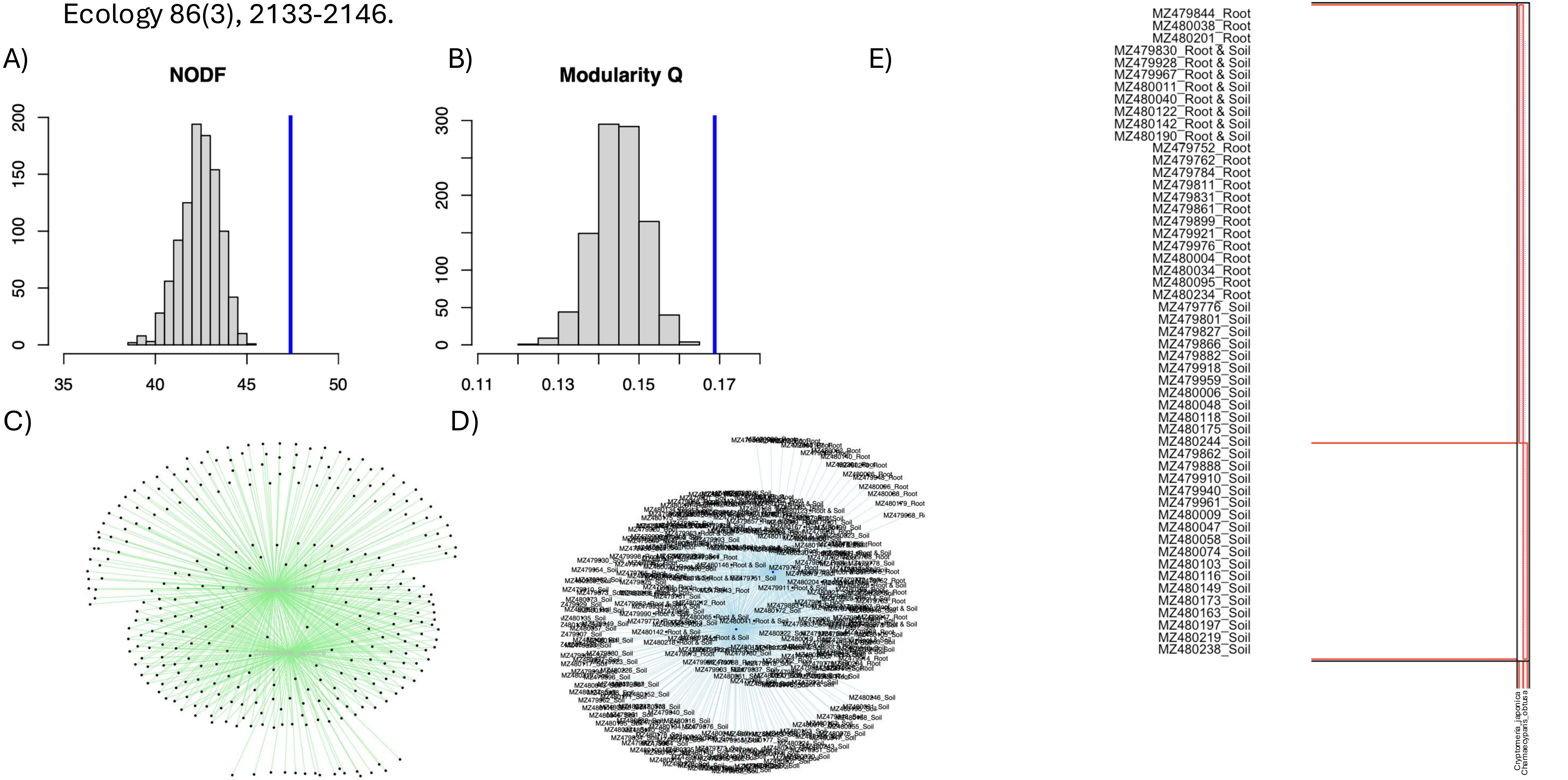

Feng, H., Wang, C., Jia, P., Gai, J. and Yang, Z. 2021. Molecular diversity of arbuscular mycorrhizal fungi associated with two alpine plant species in the Tibetan Plateau. Rhizosphere 19, 100384.

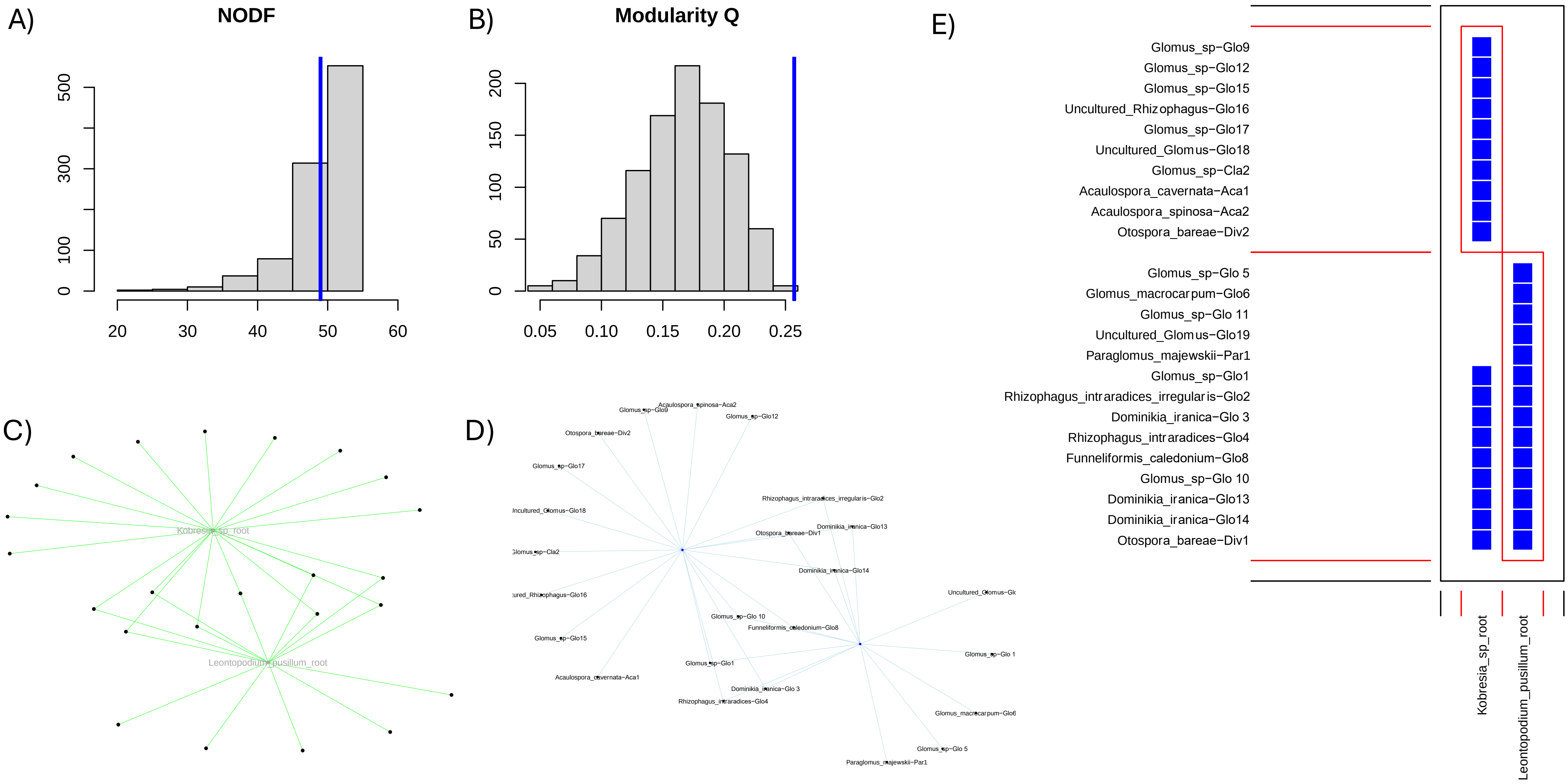

Fernández, N., Knoblochová, T., Kohout, P., Janoušková, M., Cajthaml, T., Frouz, J. and Rydlová, J. 2022. Asymmetric interaction between two mycorrhizal fungal guilds and consequences for the establishment of their host plants. Frontiers in Plant Science 13, 873204.

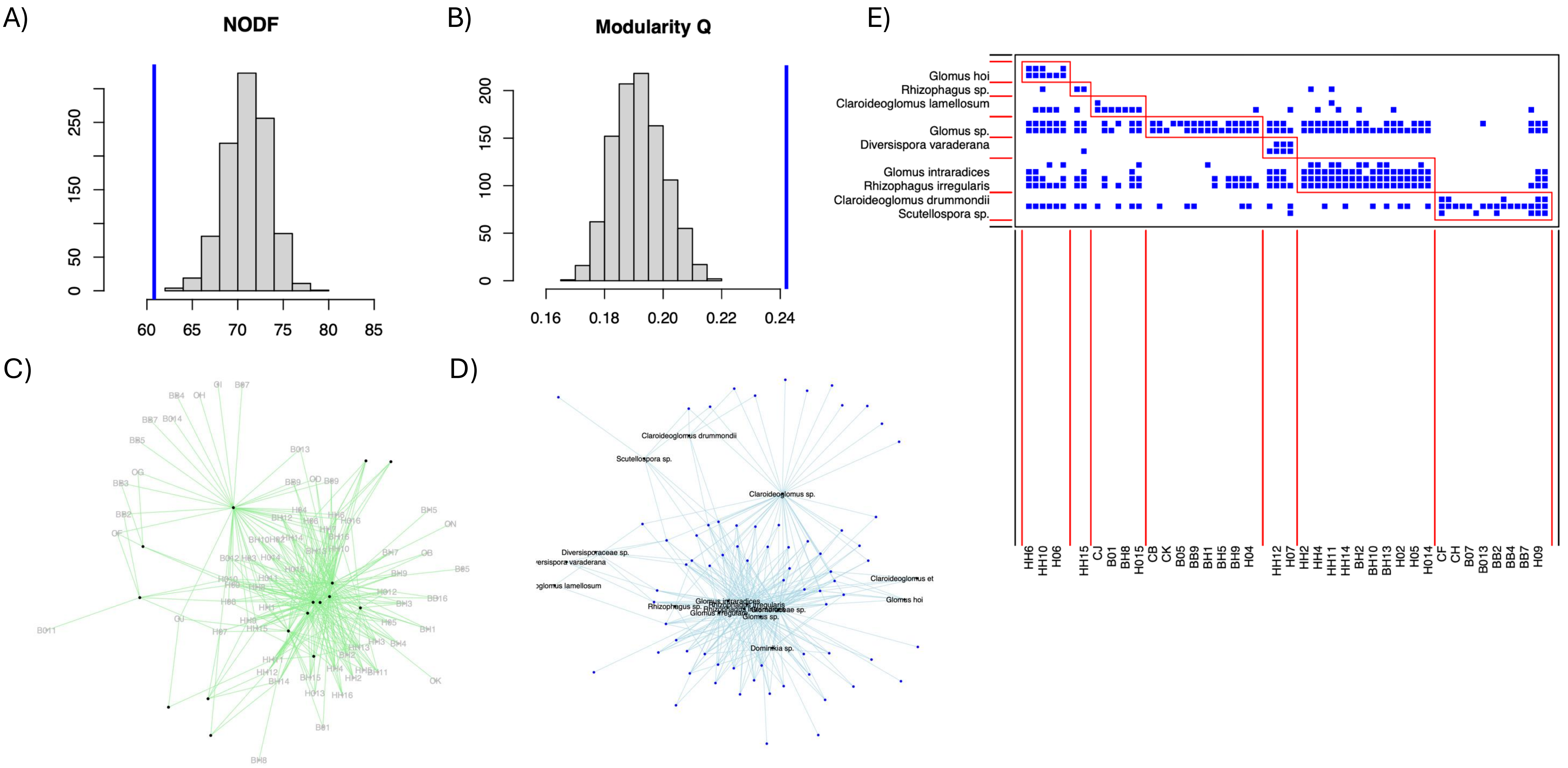

Galván, G.A., Parádi, I., Burger, K., Baar, J., Kuyper, T.W., Scholten, O.E. and Kik, C. 2009. Molecular diversity of arbuscular mycorrhizal fungi in onion roots from organic and conventional farming systems in the Netherlands. Mycorrhiza 19, 317-328.

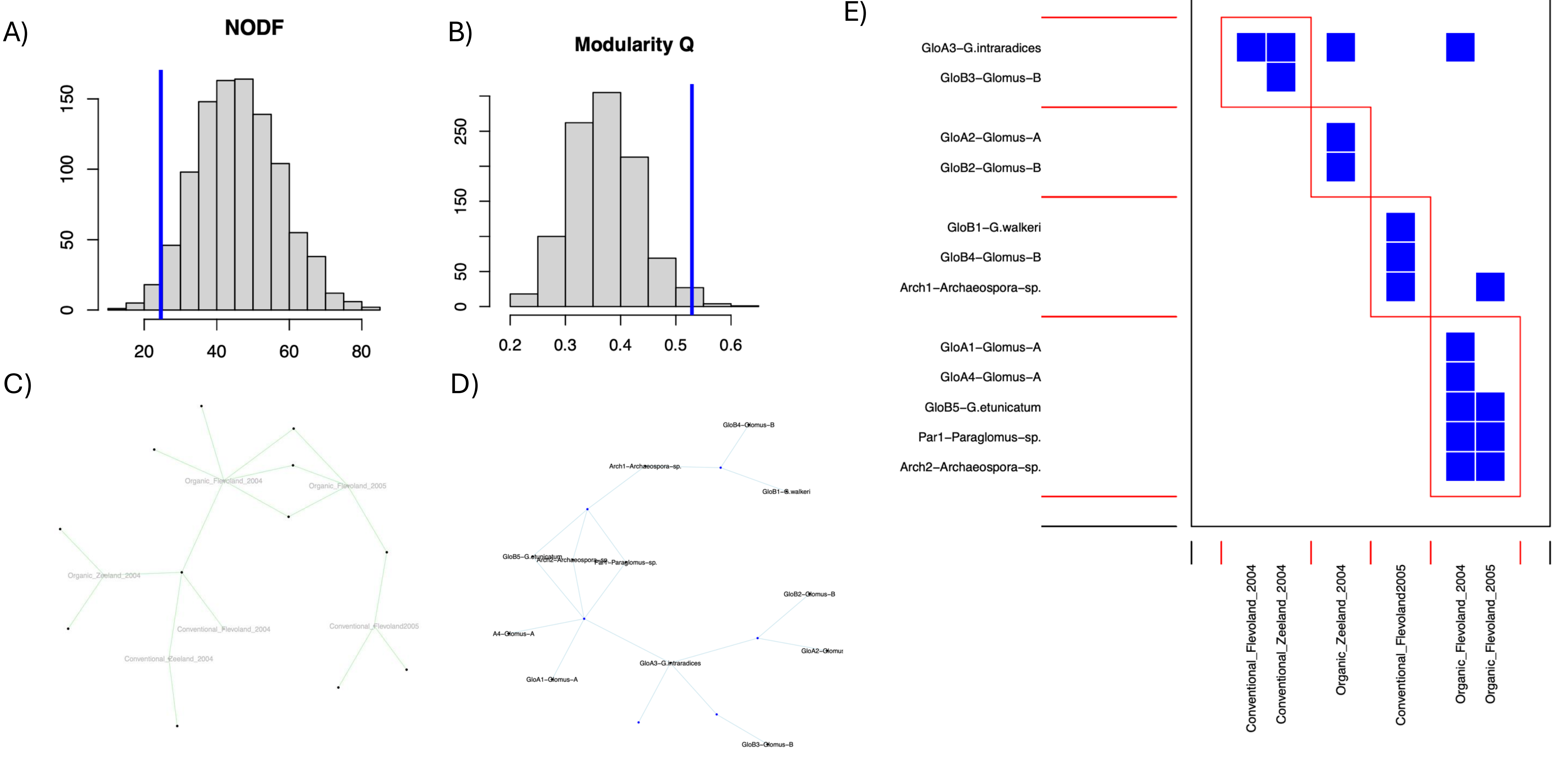

Garrido, J.L., Alcántara, J.M., López-García, Á., Ozuna, C.V., Perea, A.J., Prieto, J., Rincón, A. and Azcón-Aguilar, C. 2023. The structure and ecological function of the interactions between plants and arbuscular mycorrhizal fungi through multilayer networks. Functional Ecology 37(8), 2217-2230.

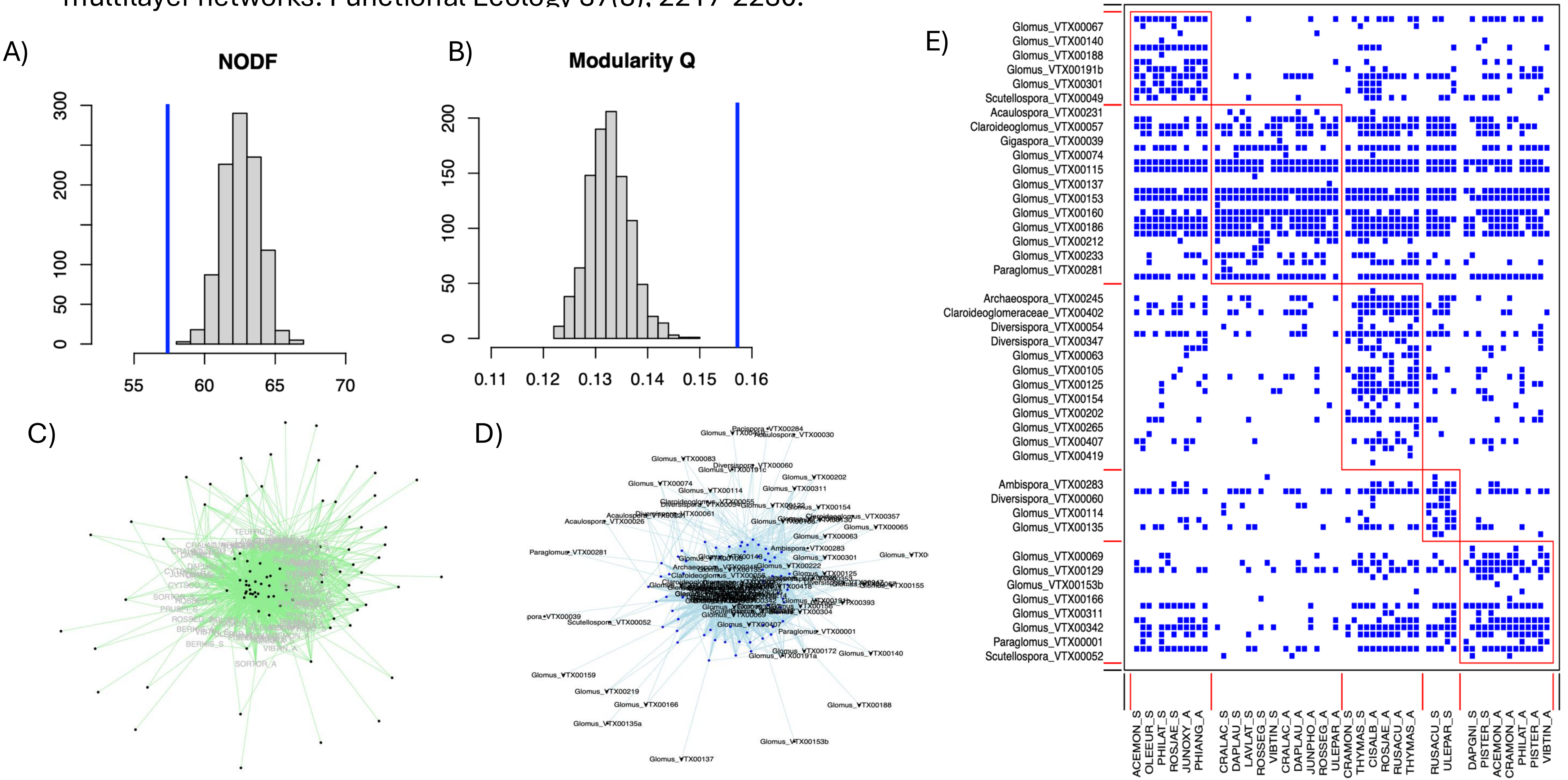

Grilli, G., Urcelay, C., Galetto, L., Davison, J., Vasar, M., Saks, Ü., Jairus, T. and Öpik, M. 2015. The composition of arbuscular mycorrhizal fungal communities in the roots of a ruderal forb is not related to the forest fragmentation process. Environmental Microbiology 17(8), 2709-2720.

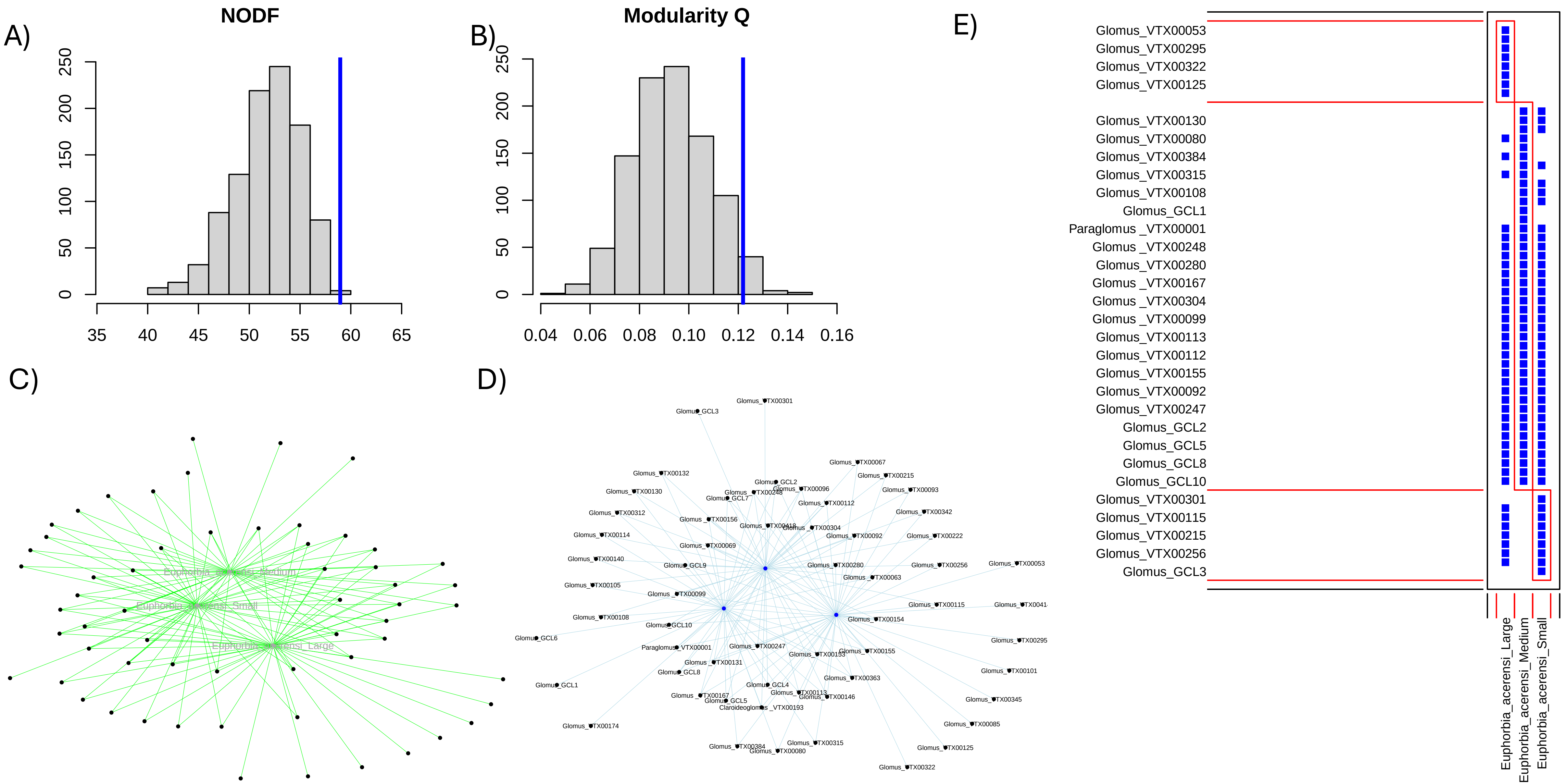

Guo, X. and Gong, J. 2014. Differential effects of abiotic factors and host plant traits on diversity and community composition of root-colonizing arbuscular mycorrhizal fungi in a salt-stressed ecosystem. Mycorrhiza 24, 79-94.

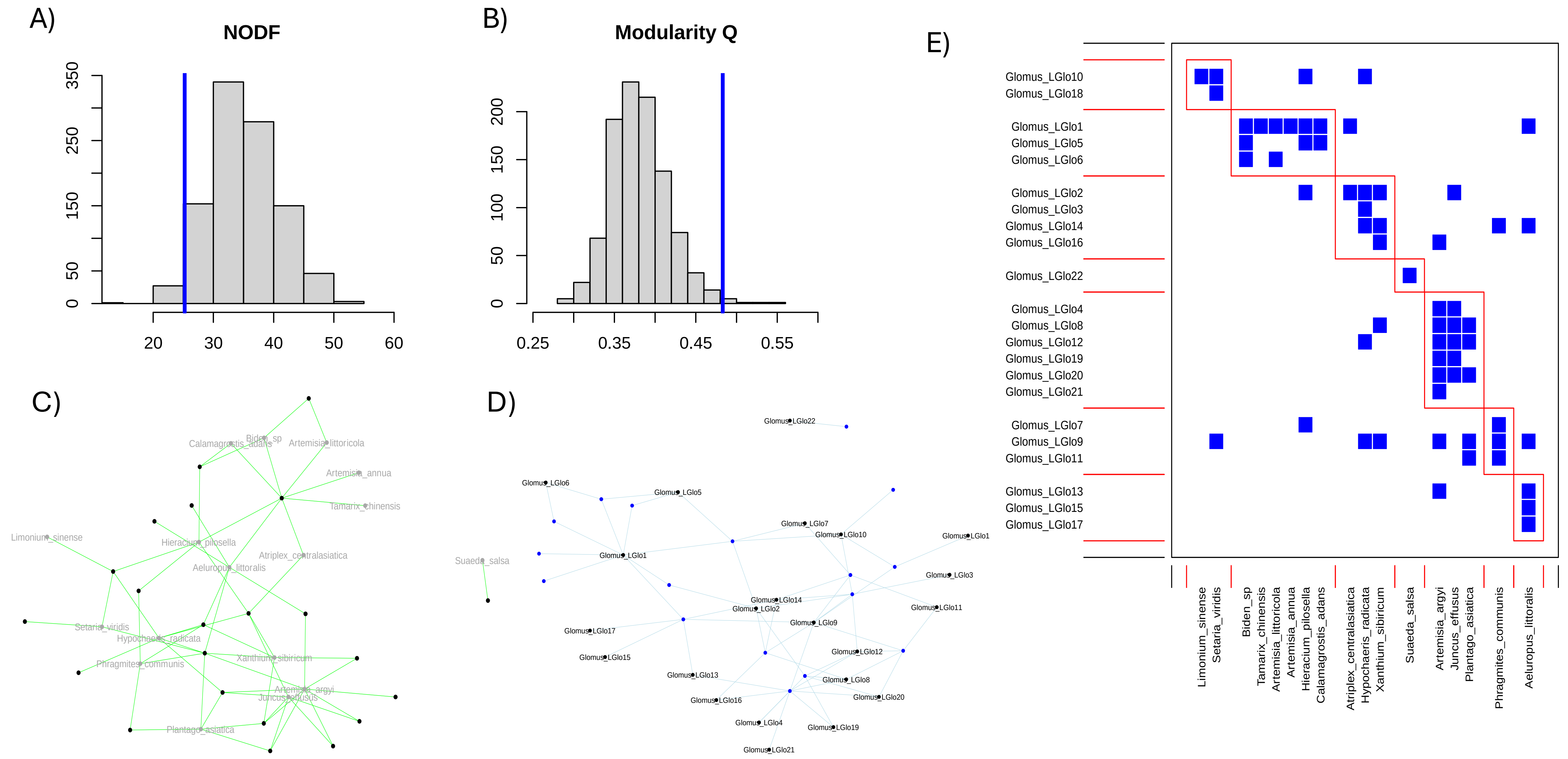

Higo, M., Tatewaki, Y., Iida, K., Yokota, K. and Isobe, K. 2020. Amplicon sequencing analysis of arbuscular mycorrhizal fungal communities colonizing maize roots in different cover cropping and tillage systems. Scientific Reports 10(1), 6039.

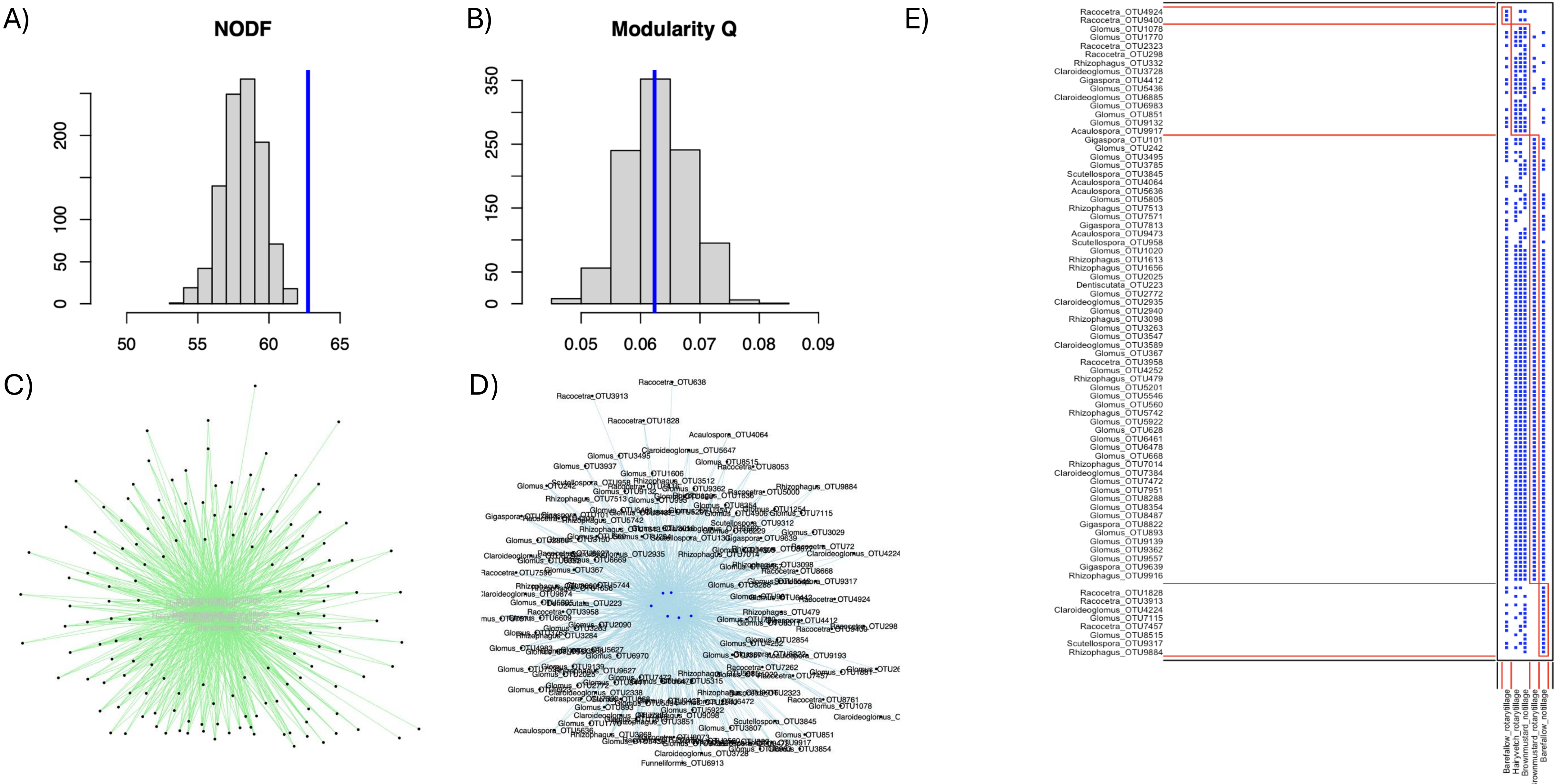

Isobe, K., Maruyama, K., Nagai, S., Higo, M., Maekawa, T., Mizonobe, G., Drijber, R.A. and Ishii, R. 2011. Arbuscular mycorrhizal fungal community structure in soybean roots: comparison between Kanagawa and Hokkaido, Japan. Advances in Microbiology 1(01), 13.

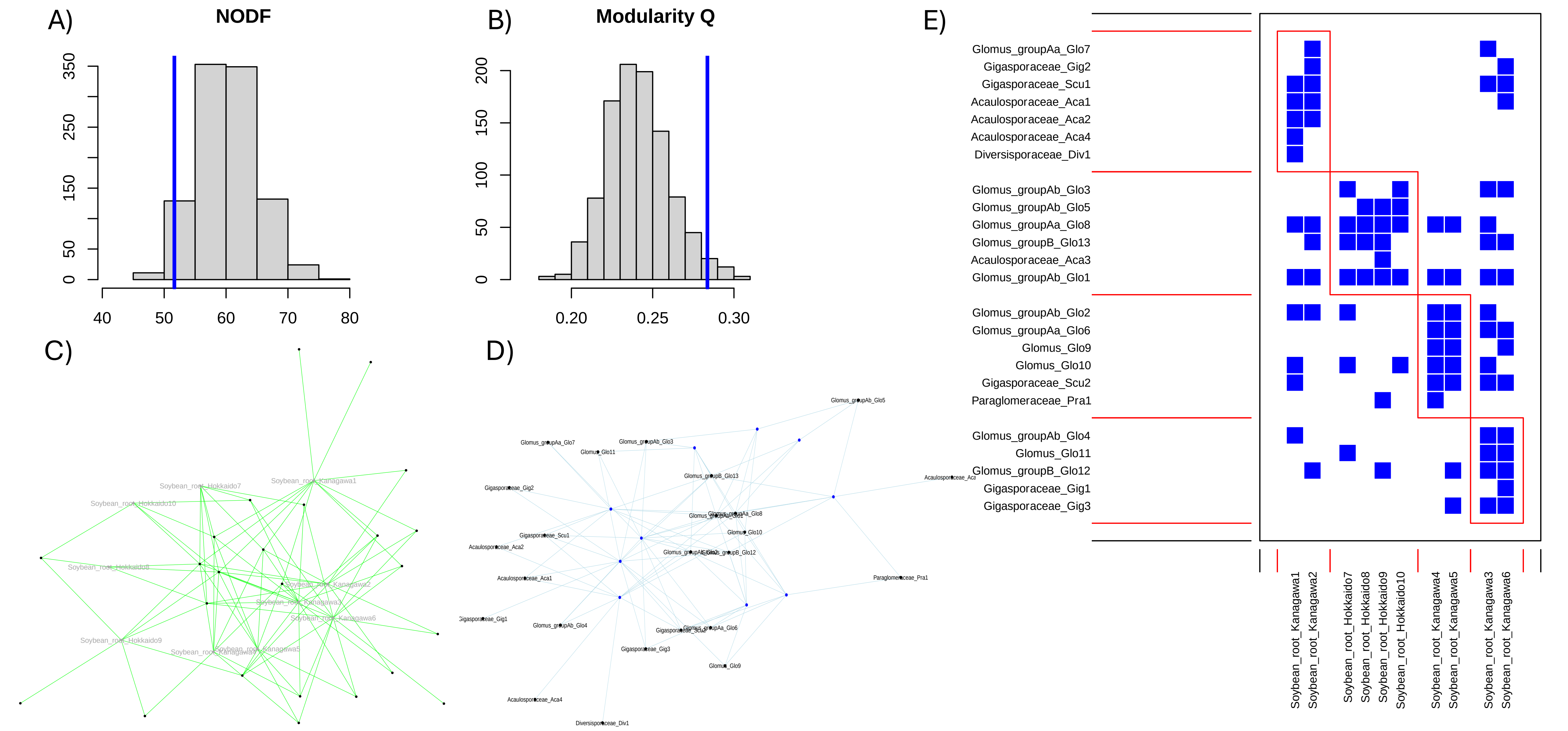

Lara-Pérez, L.A., Oros-Ortega, I., Córdova-Lara, I., Estrada-Medina, H., O'Connor-Sánchez, A., Góngora-Castillo, E. and Sáenz-Carbonell, L. 2020. Seasonal shifts of arbuscular mycorrhizal fungi in Cocos nucifera roots in Yucatan, Mexico. Mycorrhiza 30, 269-283.

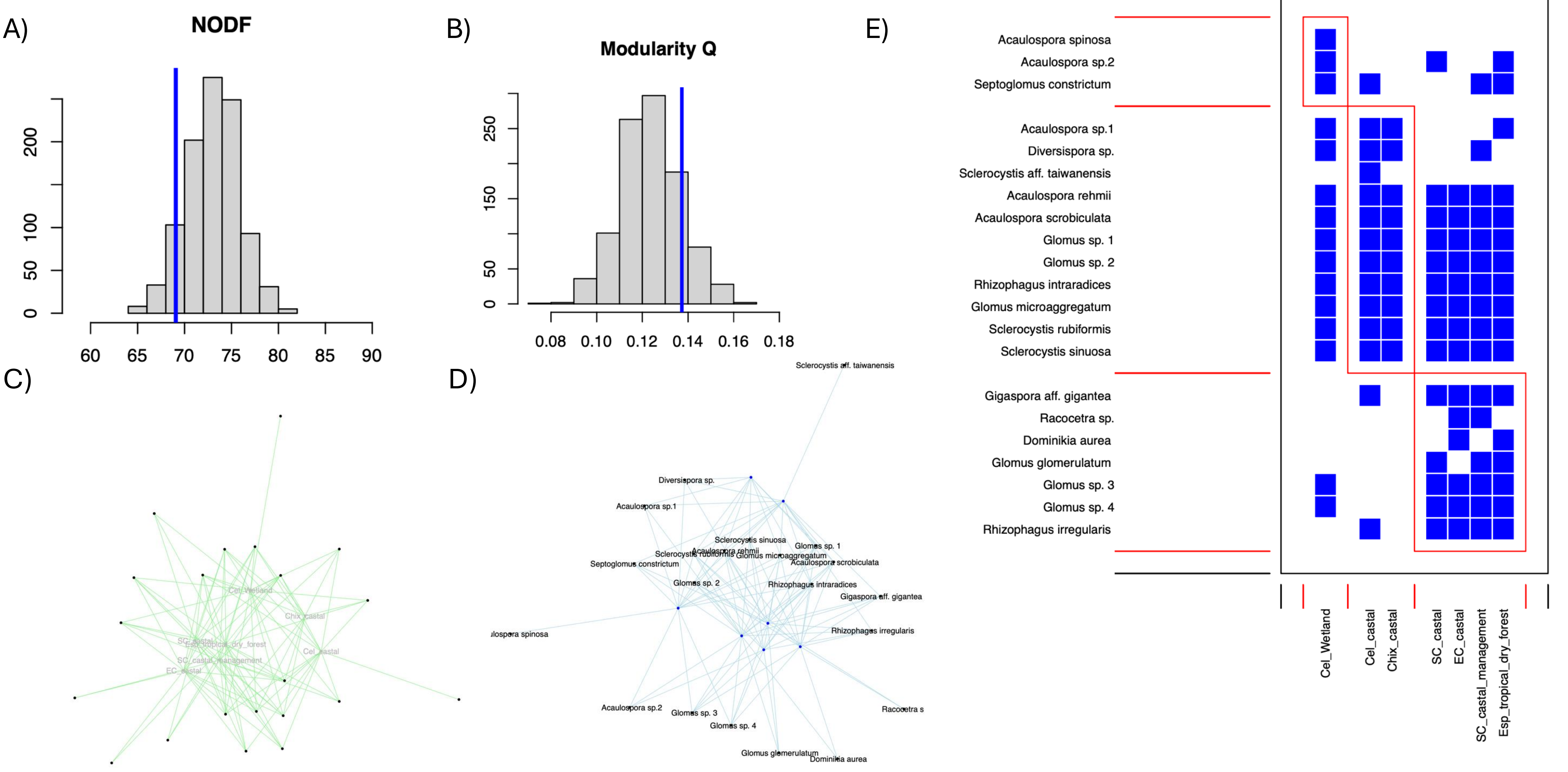

Li, X., Gai, J., Cai, X., Li, X., Christie, P., Zhang, F. and Zhang, J. 2014. Molecular diversity of arbuscular mycorrhizal fungi associated with two co-occurring perennial plant species on a Tibetan altitudinal gradient. Mycorrhiza 24, 95-107.

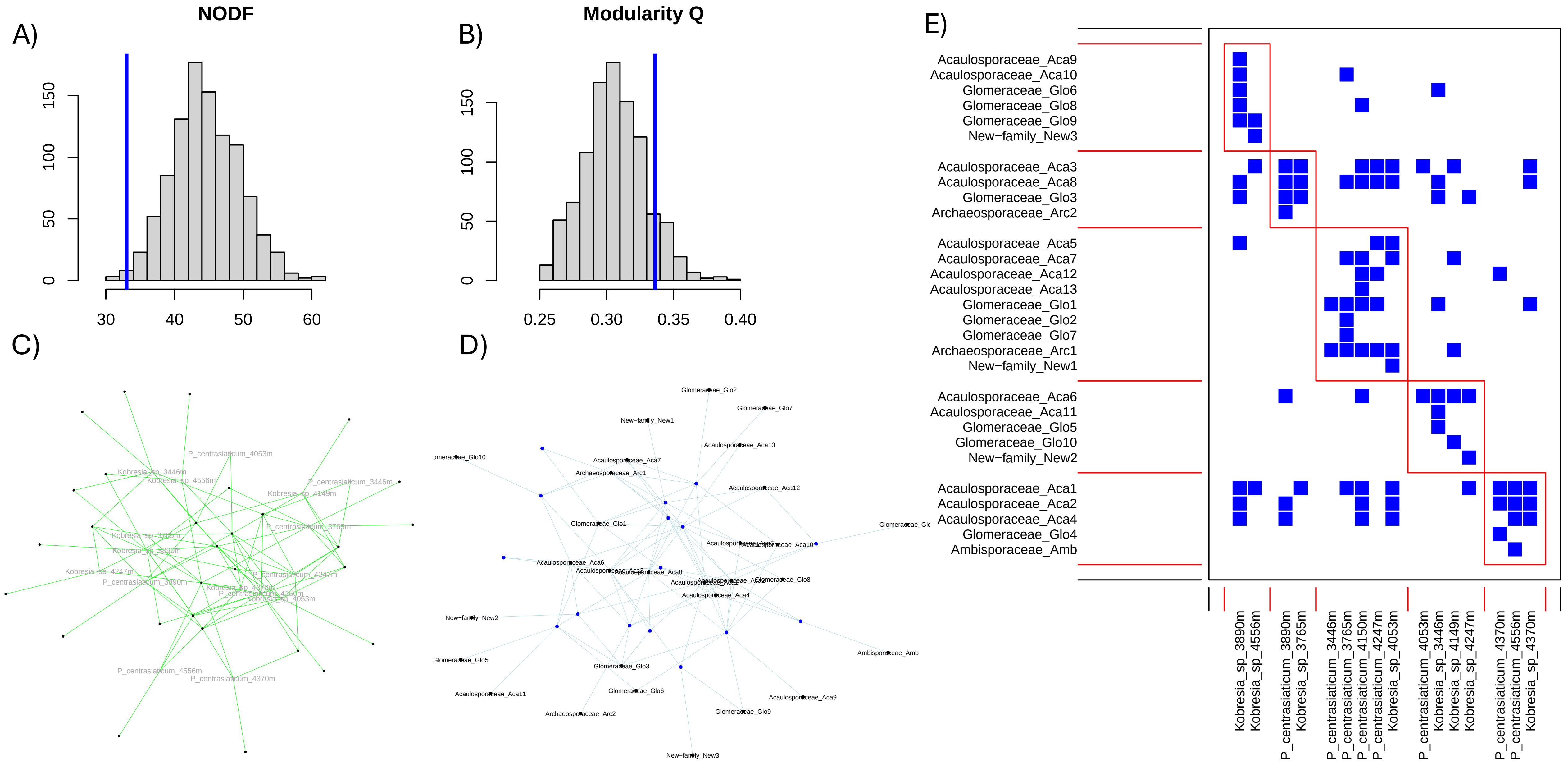

Liu, Y., He, J., Shi, G., An, L., Öpik, M. and Feng, H. 2011. Diverse communities of arbuscular mycorrhizal fungi inhabit sites with very high altitude in Tibet Plateau. FEMS Microbiology Ecology 78(2), 355-365.

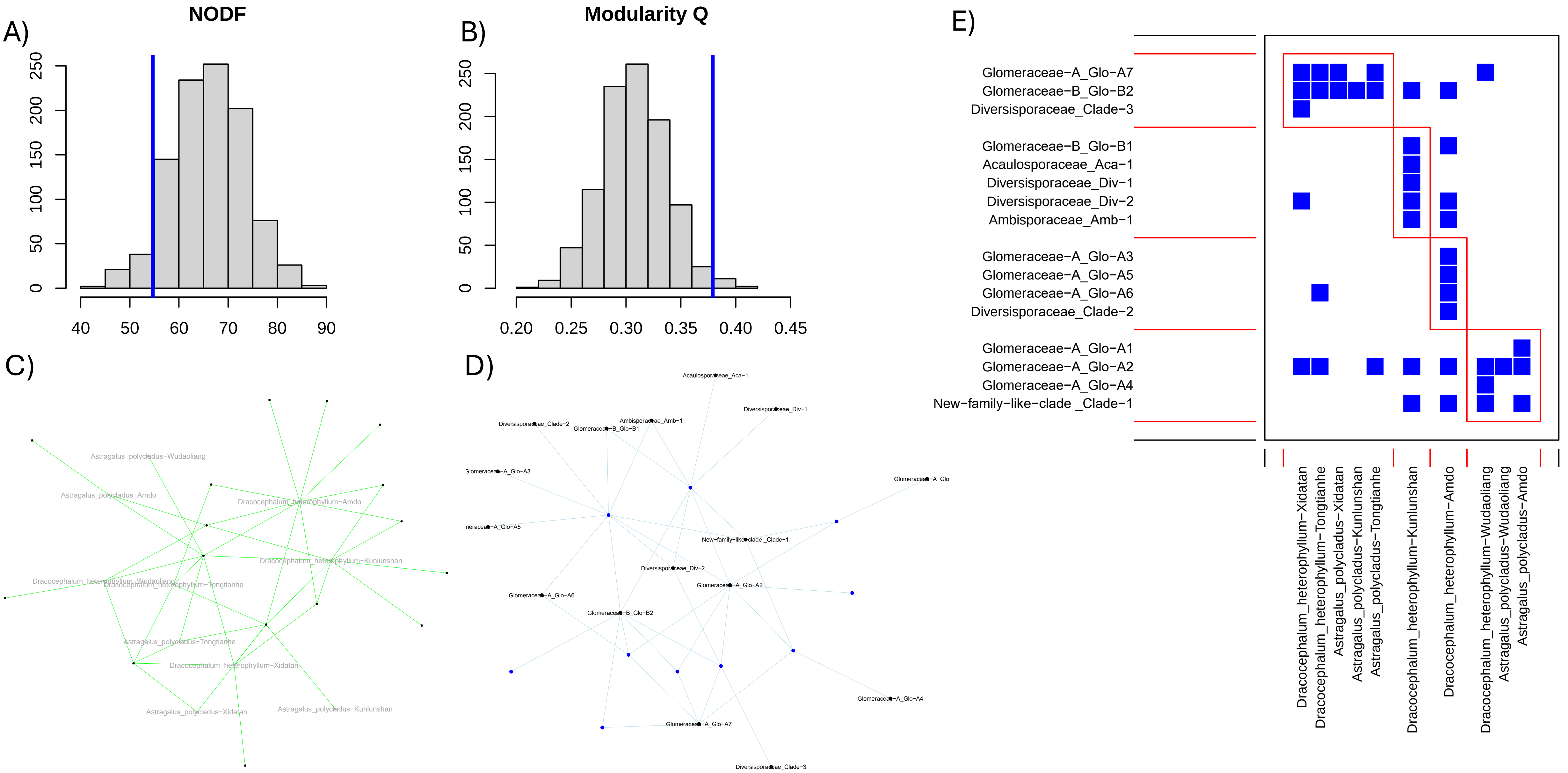

Moora, M., Berger, S., Davison, J., Öpik, M., Bommarco, R., Bruelheide, H., Kühn, I., Kunin, W.E., Metsis, M. and Rortais, A. 2011. Alien plants associate with widespread generalist arbuscular mycorrhizal fungal taxa: evidence from a continental-scale study using massively parallel 454 sequencing. Journal of Biogeography 38(7), 1305-1317.

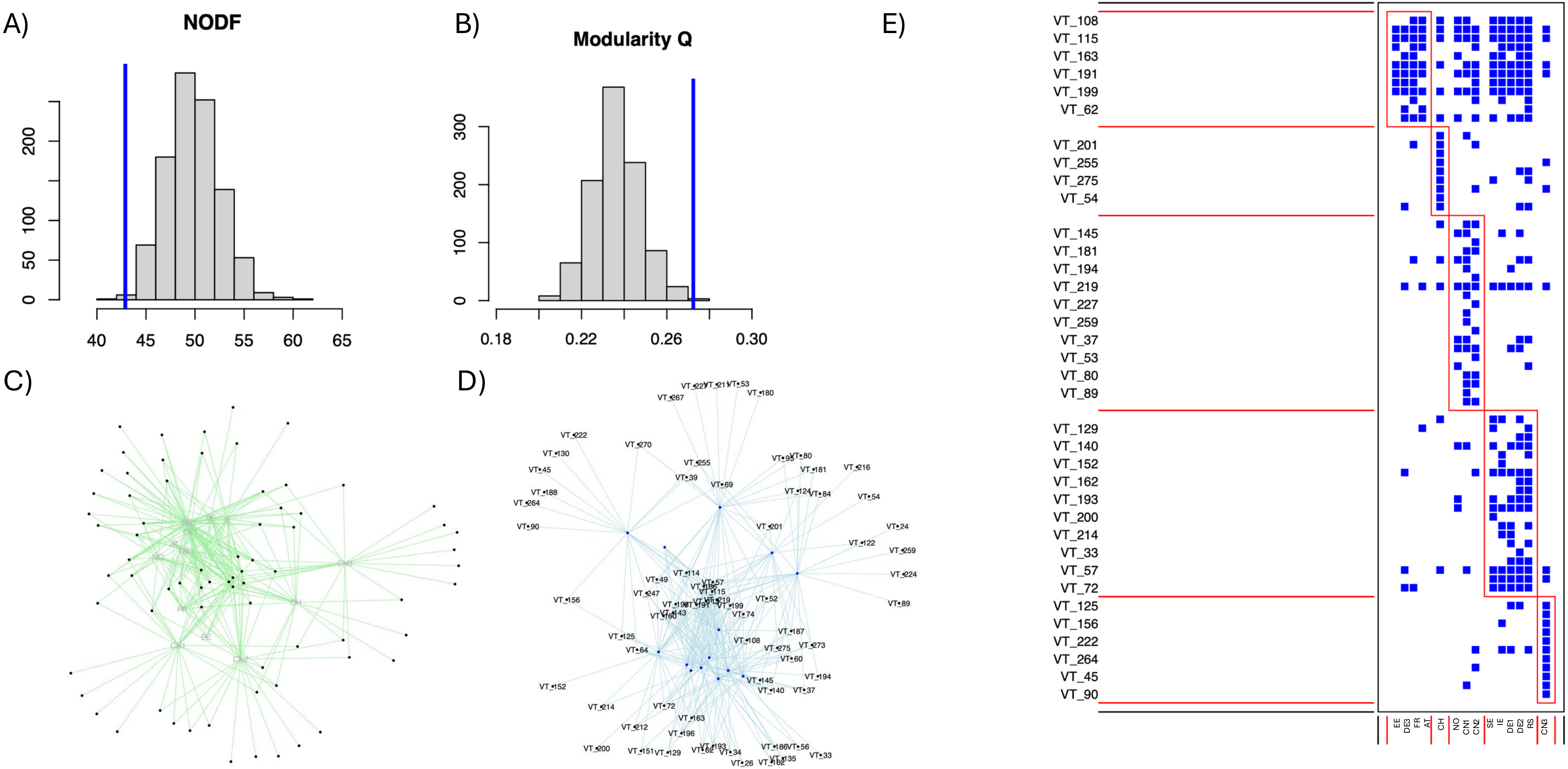

Moora, M., Öpik, M., Davison, J., Jairus, T., Vasar, M., Zobel, M. and Eckstein, R.L. 2016. AM fungal communities inhabiting the roots of submerged aquatic plant Lobelia dortmanna are diverse and include a high proportion of novel taxa. Mycorrhiza 26, 735–745.

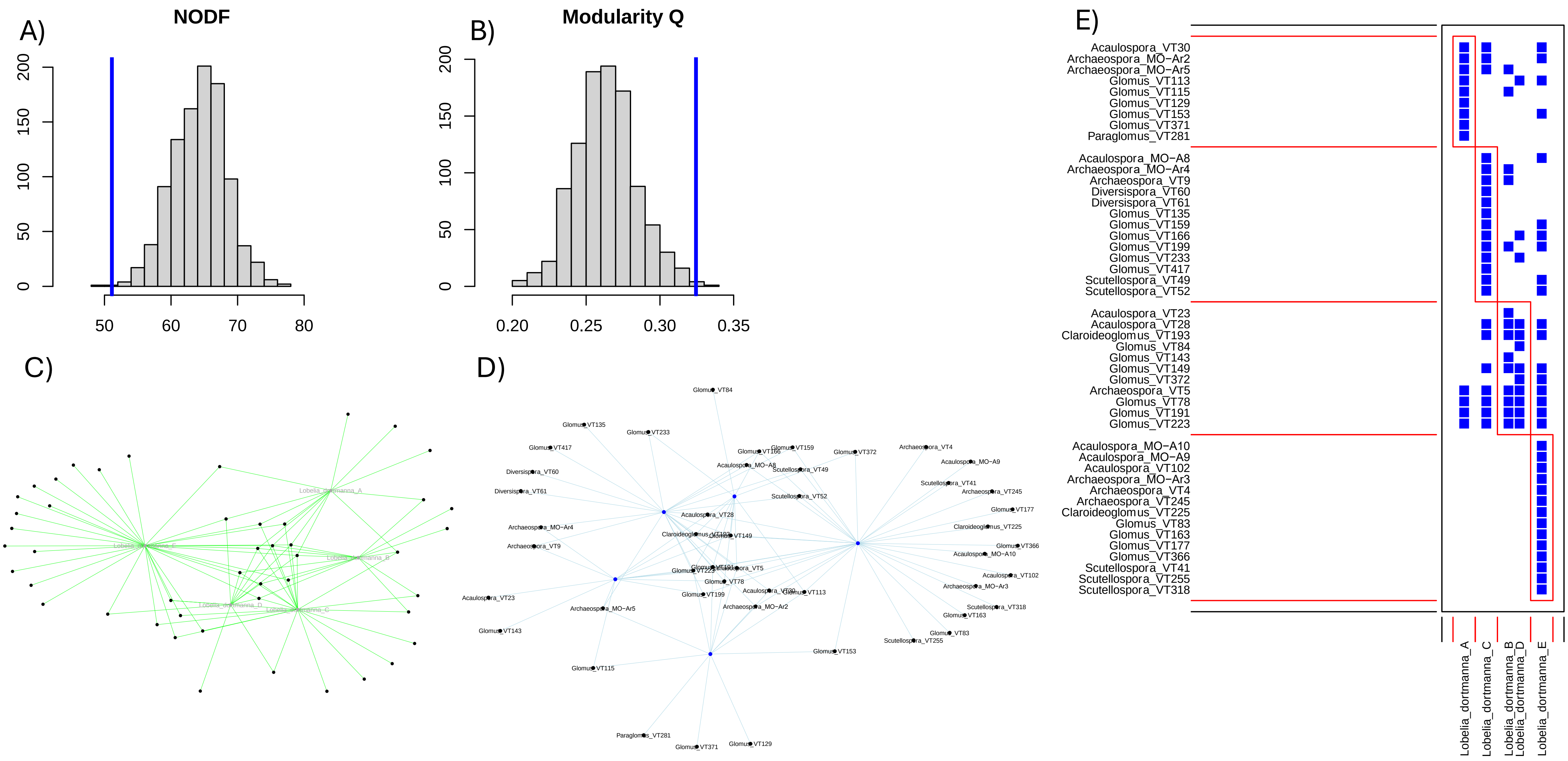

Muneer, M., WANG, Z., Jing, Z., Zhou, X., Wang, P., Li, L. and Ji, B. 2019. Low host specificity of arbuscular mycorrhizal fungi associated with dominant steppe plants in Inner Mongolia. Applied Ecology & Environmental Research 17(5).

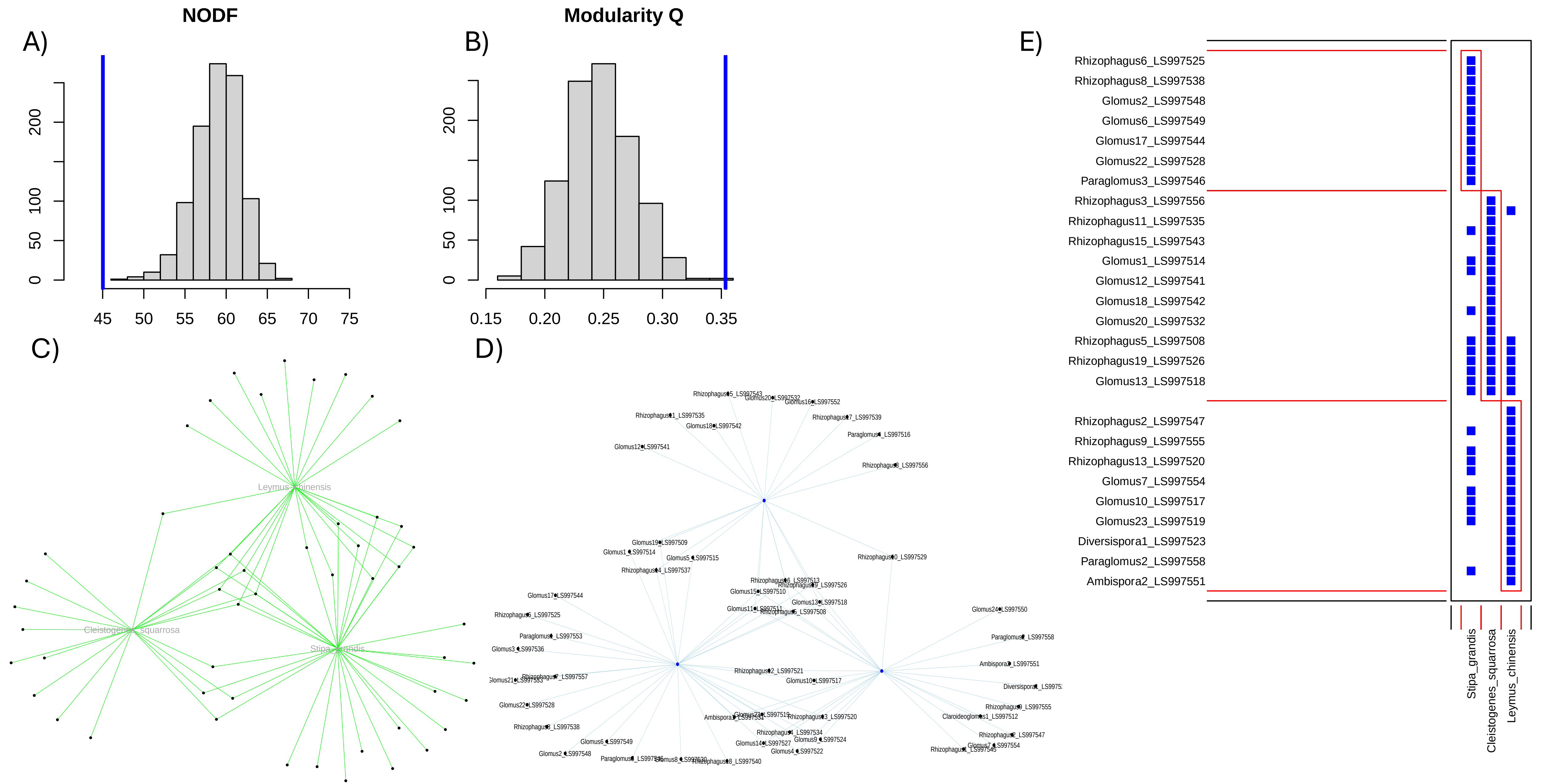

Öpik, M., Metsis, M., Daniell, T., Zobel, M. and Moora, M. 2009. Large-scale parallel 454 sequencing reveals host ecological group specificity of arbuscular mycorrhizal fungi in a boreonemoral forest. New Phytologist 184(2), 424-437

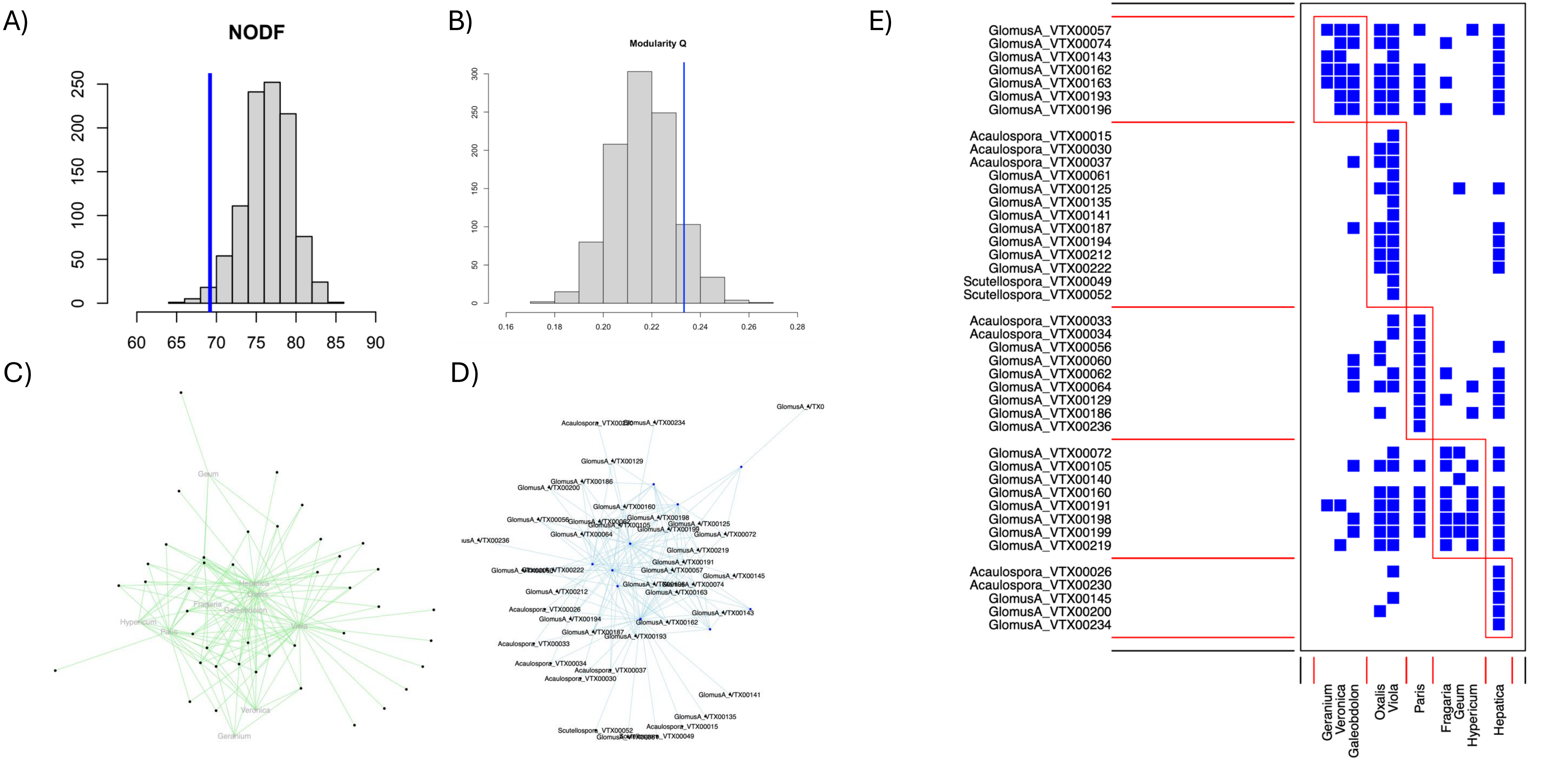

Polme, S., Öpik, M., Moora, M., Zobel, M., Kohout, P., Oja, J., Koljalg, U. and Tedersoo, L. 2016. Arbuscular mycorrhizal fungi associating with roots of Alnus and Rubus in Europe and the Middle East. Fungal ecology 24, 27-34.

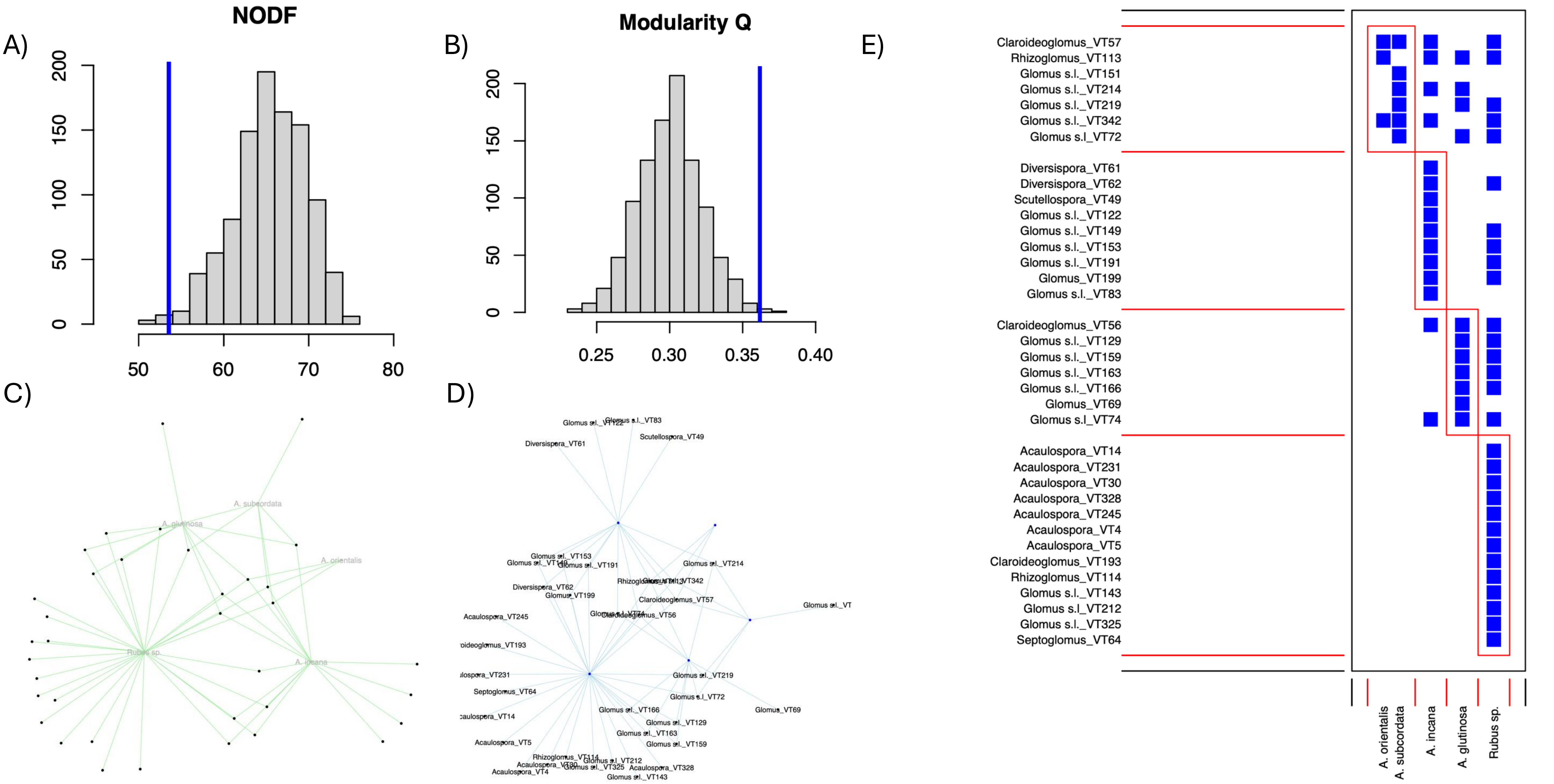

Sánchez-Castro, I., Ferrol, N. and Barea, J. 2012. Analyzing the community composition of arbuscular mycorrhizal fungi colonizing the roots of representative shrubland species in a Mediterranean ecosystem. Journal of Arid Environments 80, 1-9.

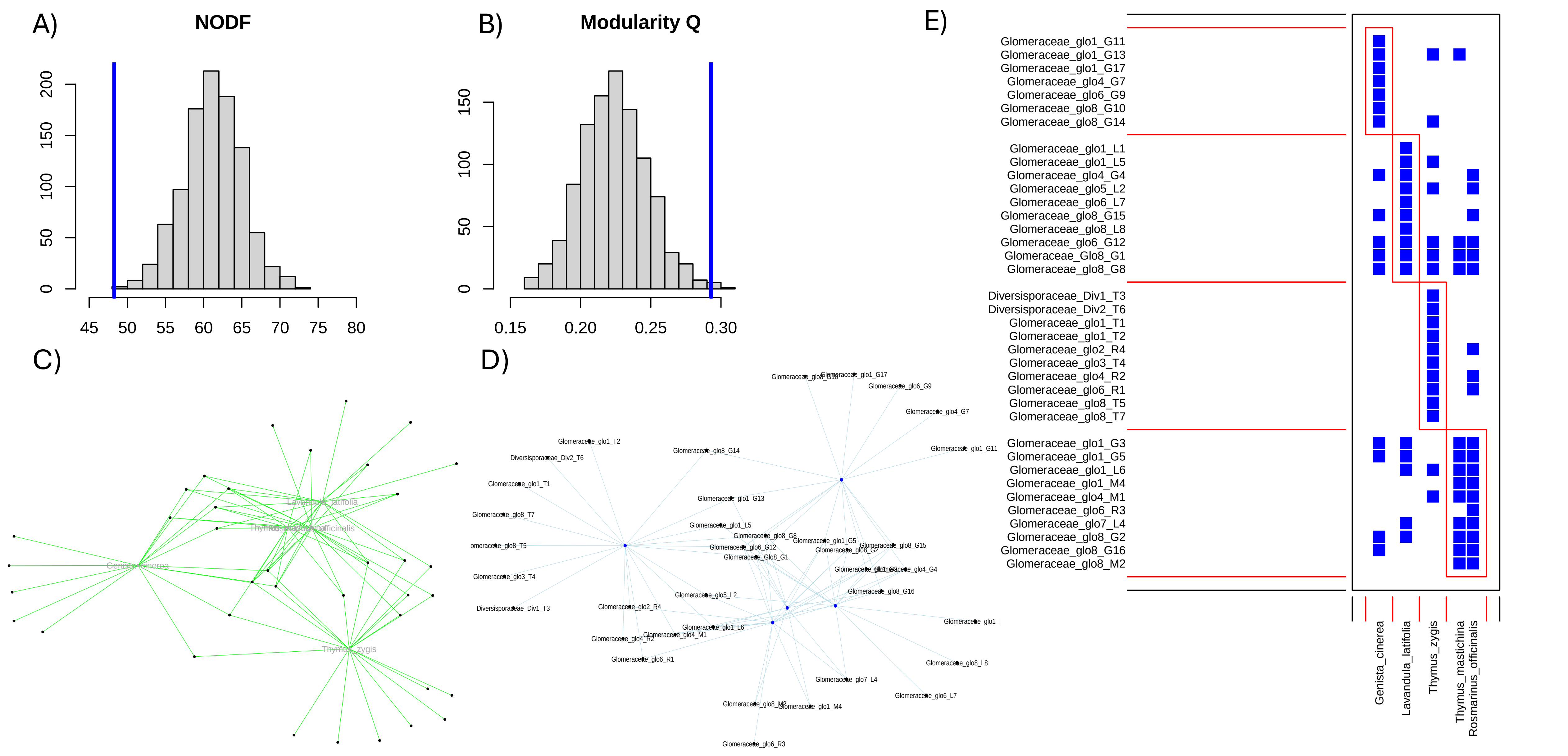

Varela-Cervero, S., Vasar, M., Davison, J., Barea, J.M., Öpik, M. and Azcón-Aguilar, C. 2015. The composition of arbuscular mycorrhizal fungal communities differs among the roots, spores and extraradical mycelia associated with five Mediterranean plant species. Environmental microbiology 17(8), 2882-2895.

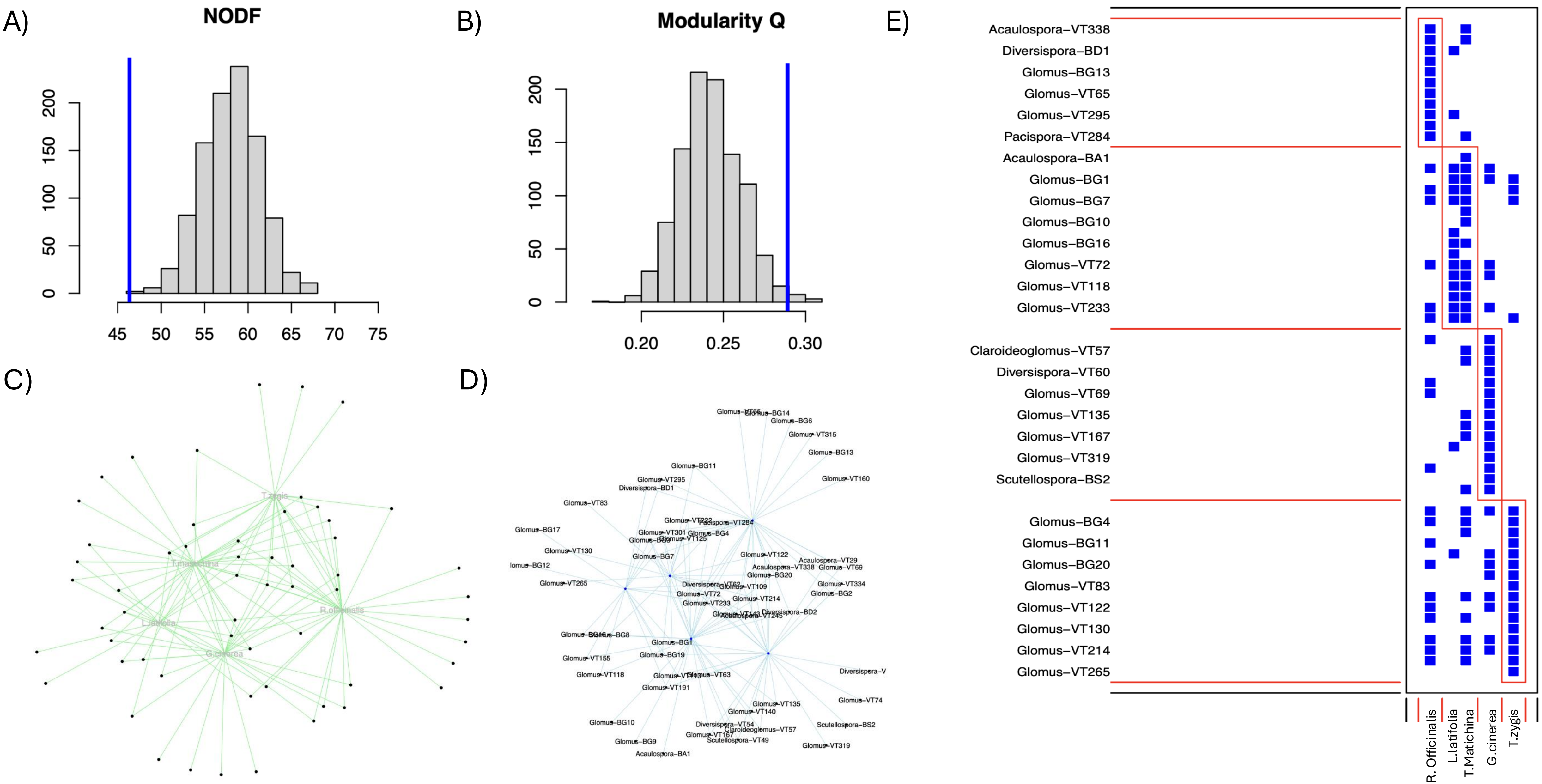

Sasvári, Z., Hornok, L. and Posta, K. 2011. The community structure of arbuscular mycorrhizal fungi in roots of maize grown in a 50-year monoculture. Biology and Fertility of Soils 47, 167-176.

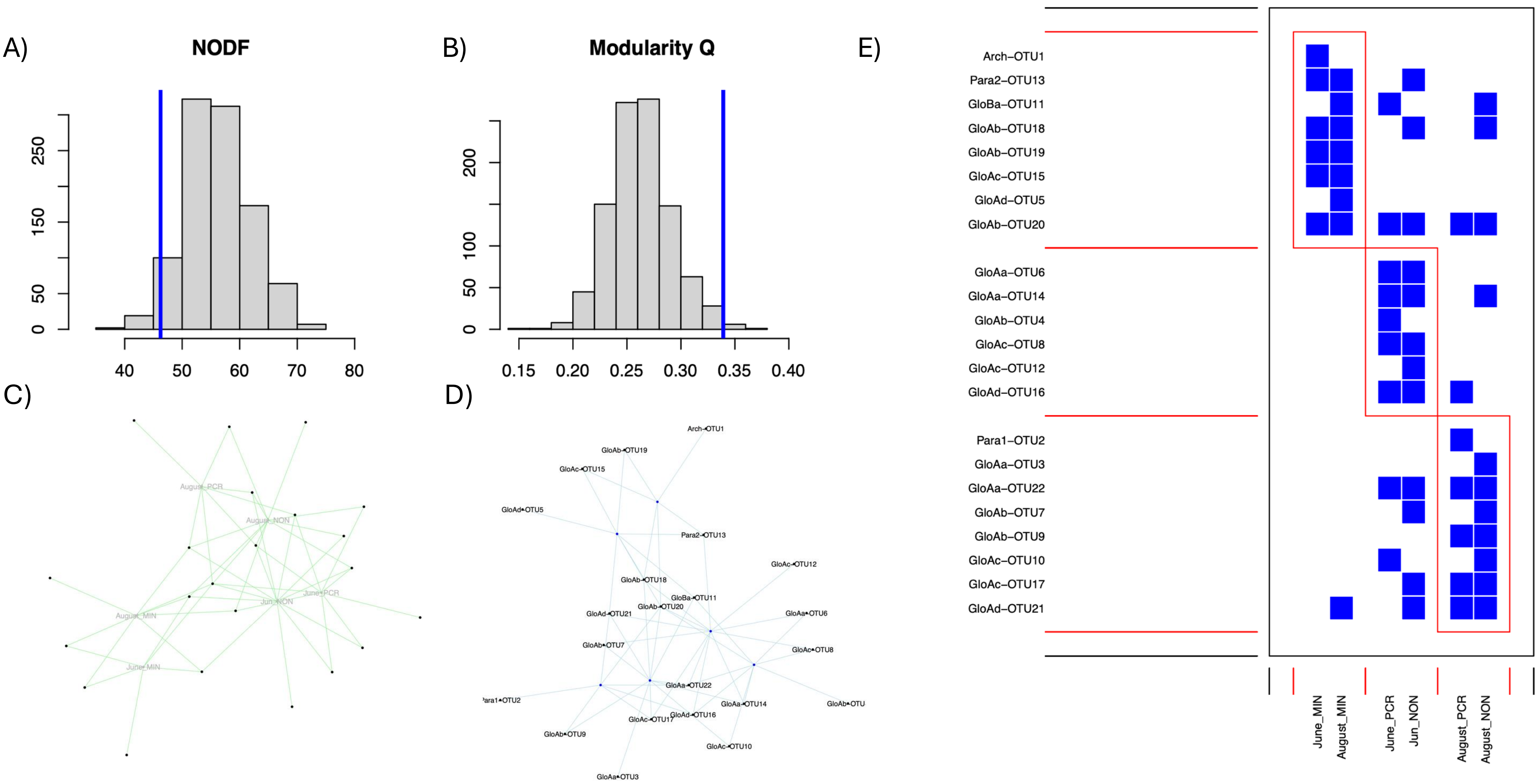

Tchabi, A., Burger, S., Coyne, D., Hountondji, F., Lawouin, L., Wiemken, A. and Oehl, F. 2009. Promiscuous arbuscular mycorrhizal symbiosis of yam (Dioscorea spp.), a key staple crop in West Africa. Mycorrhiza 19, 375-392.

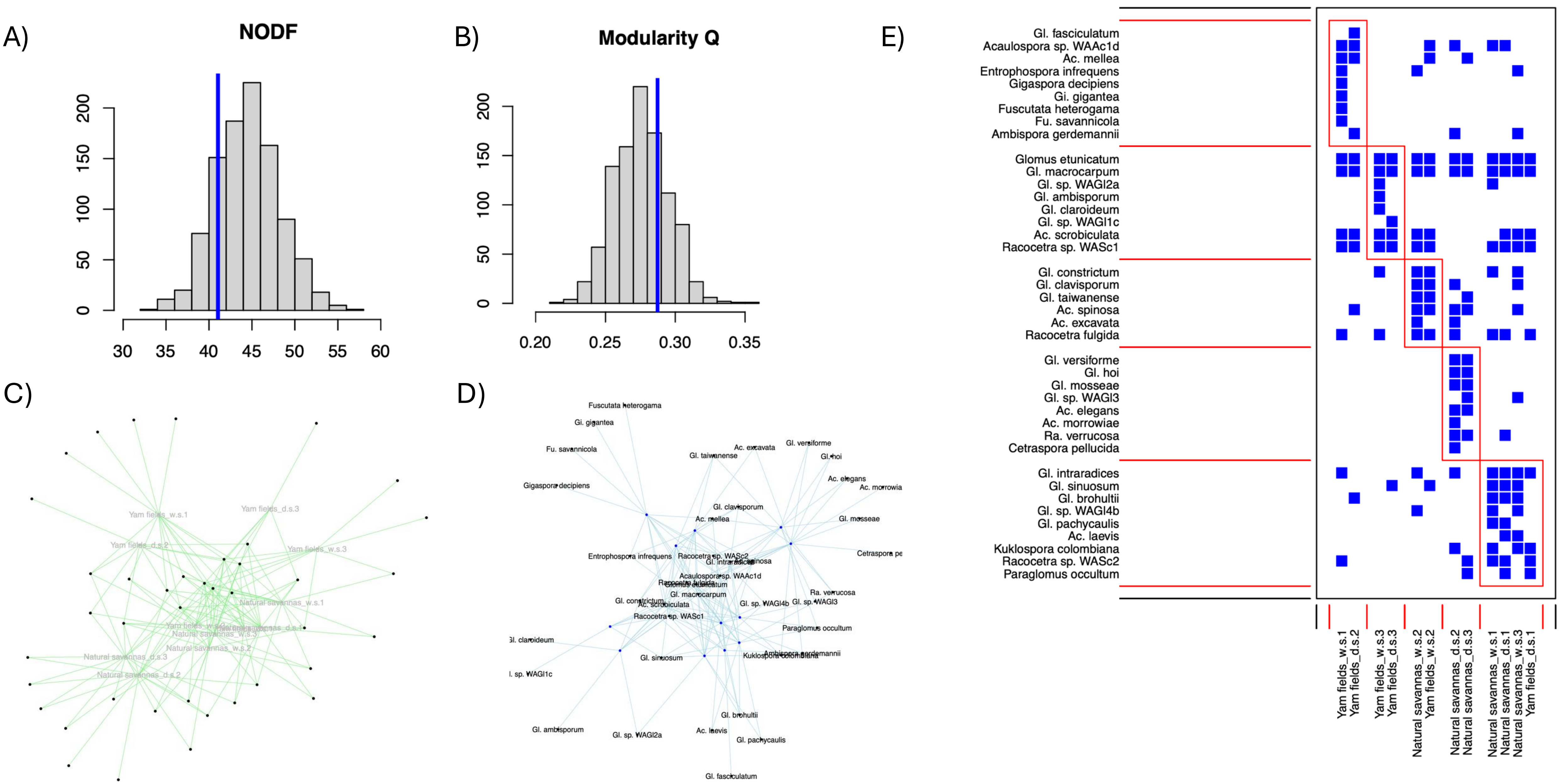

Merckx, V.S., Janssens, S.B., Hynson, N.A., Specht, C.D., Bruns, T.D. and Smets, E.F. 2012. Mycoheterotrophic interactions are not limited to a narrow phylogenetic range of arbuscular mycorrhizal fungi. Molecular Ecology 21(6), 1524-1532.

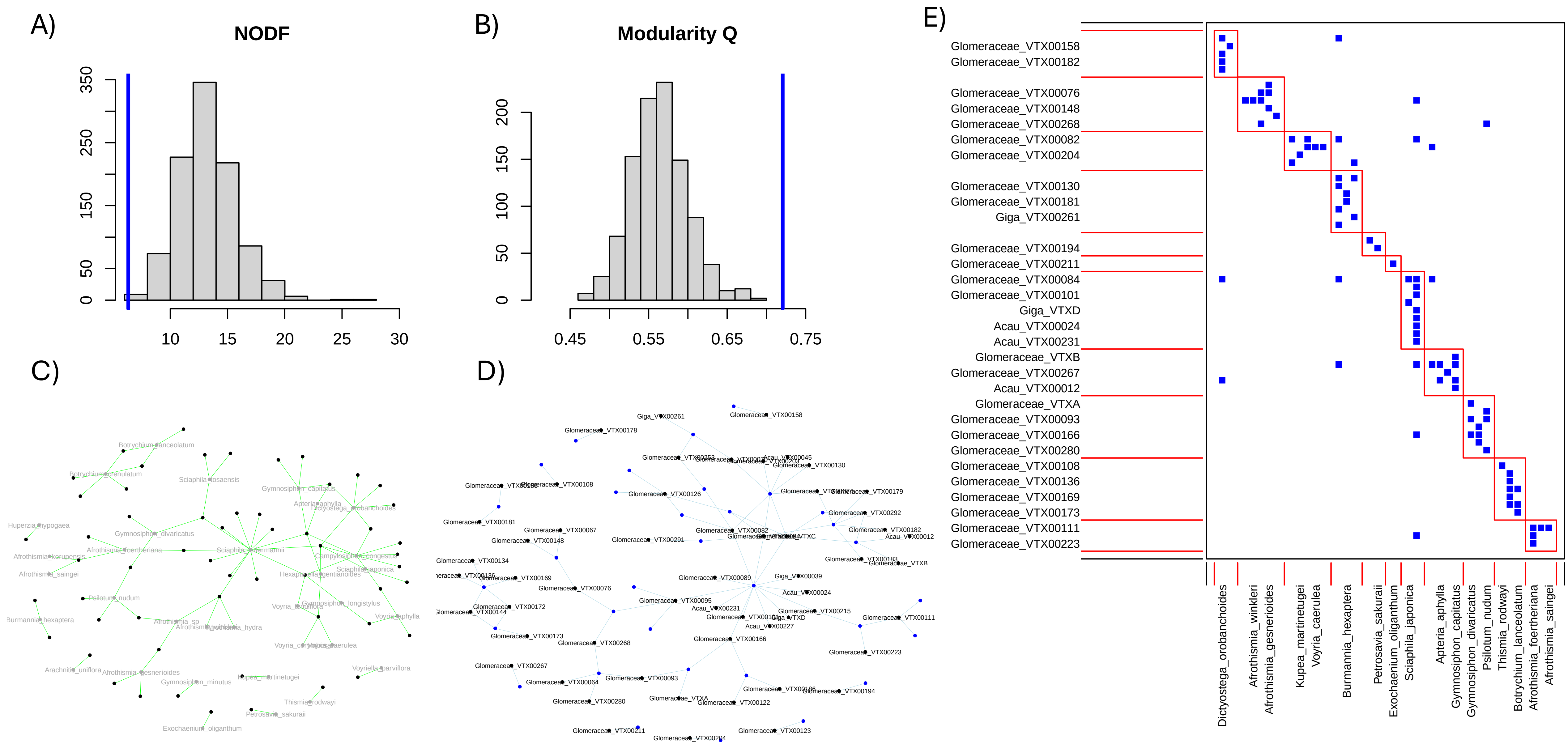

Wang, K., He, X., Xie, L. and Zhao, L. 2018. Arbuscular mycorrhizal fungal community structure and diversity are affected by host plant species and soil depth in the Mu Us Desert, northwest China. Arid Land Research and Management 32(2), 198-211.

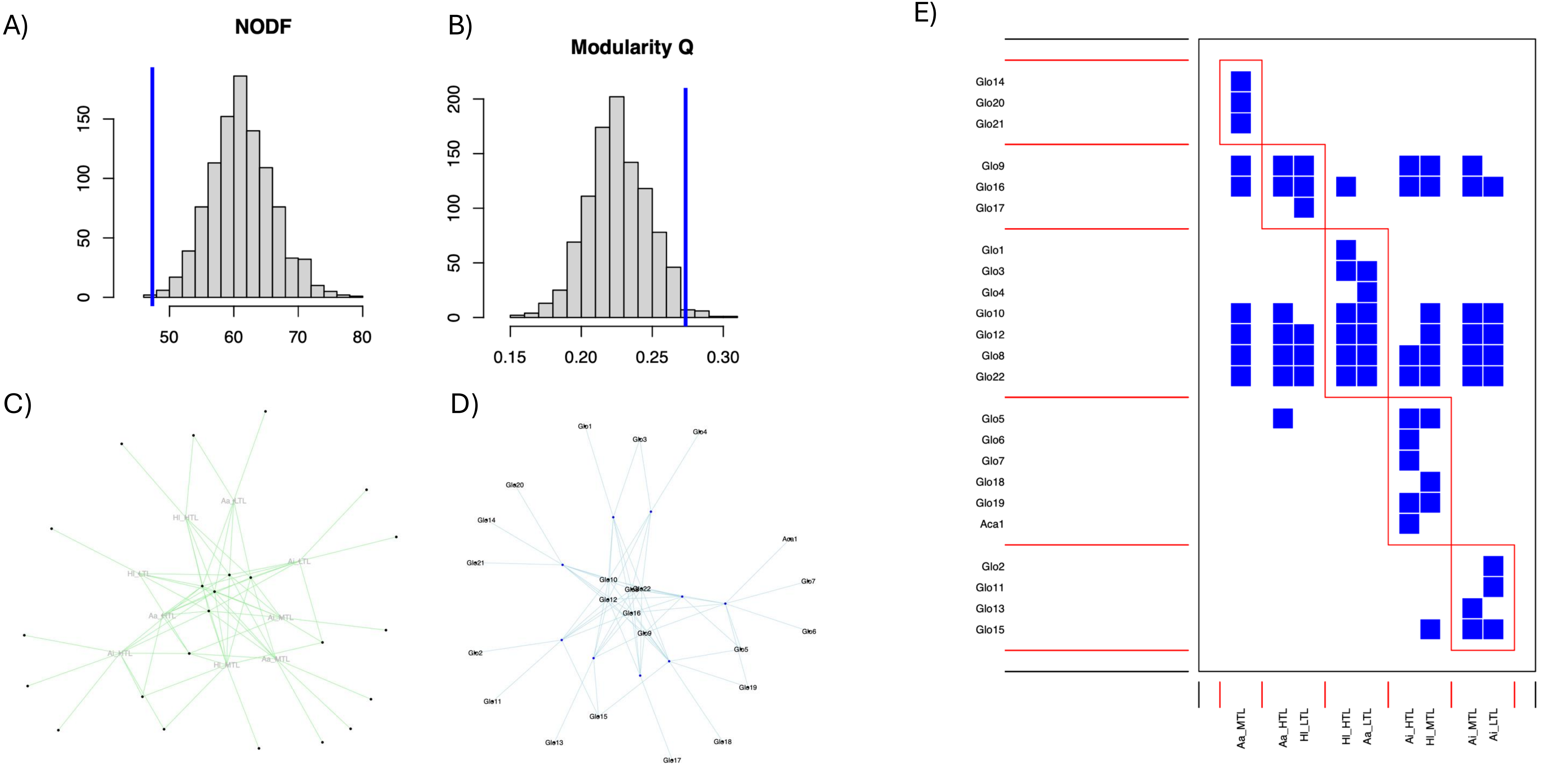

Wang, J., Gao, X., Wang, J., Song, J., Zhu, Z., Zhao, J., Liu, M. and Wang, Z. 2024. Host plants directly determine the α diversity of rhizosphere arbuscular mycorrhizal fungal communities in the National Tropical Fruit Tree Field Genebank. Chemical and Biological Technologies in Agriculture 11(1), 20.

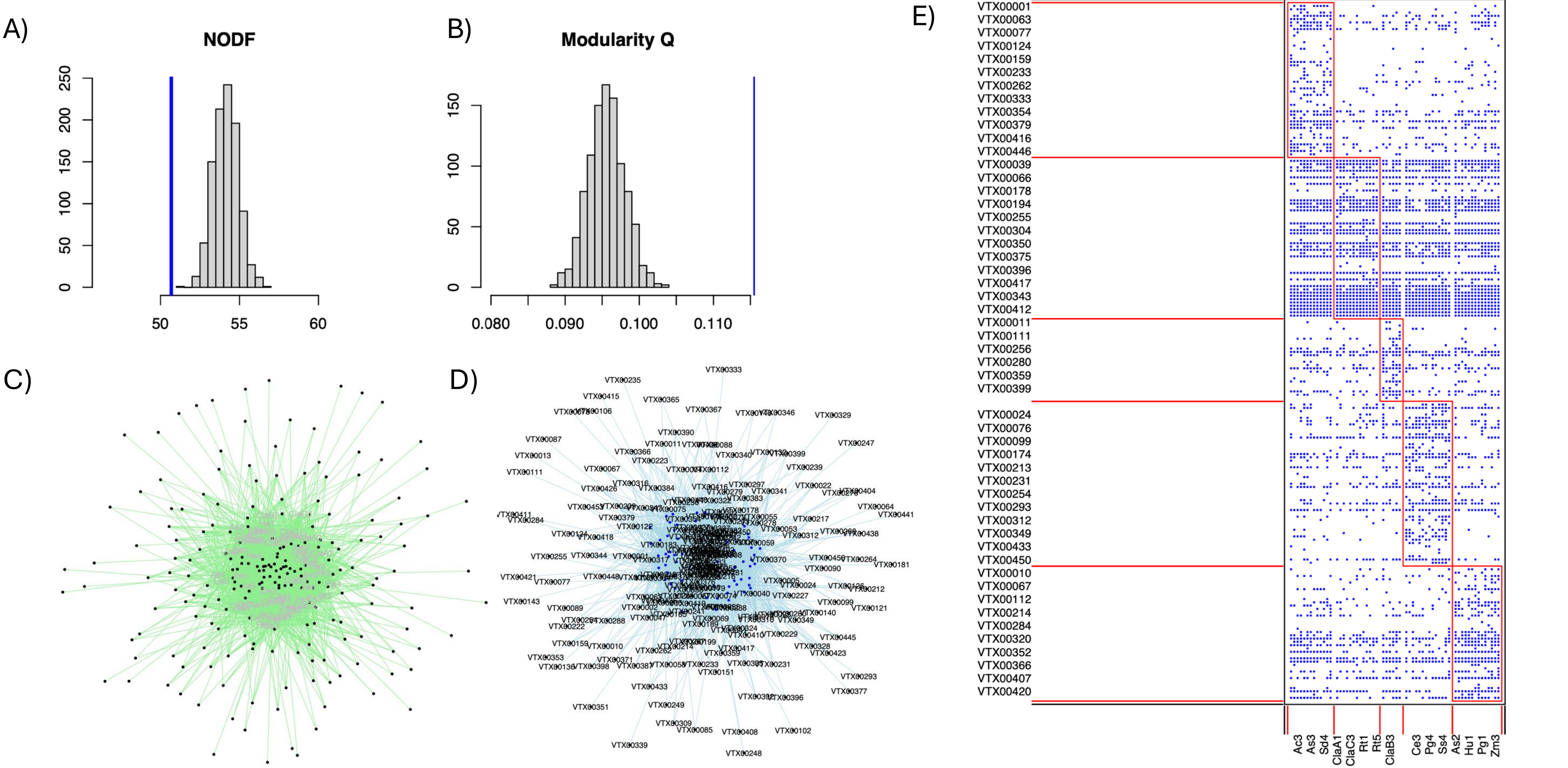

Wilde, P., Manal, A., Stodden, M., Sieverding, E., Hildebrandt, U. and Bothe, H. 2009. Biodiversity of arbuscular mycorrhizal fungi in roots and soils of two salt marshes. Environmental Microbiology 11(6), 1548-1561.

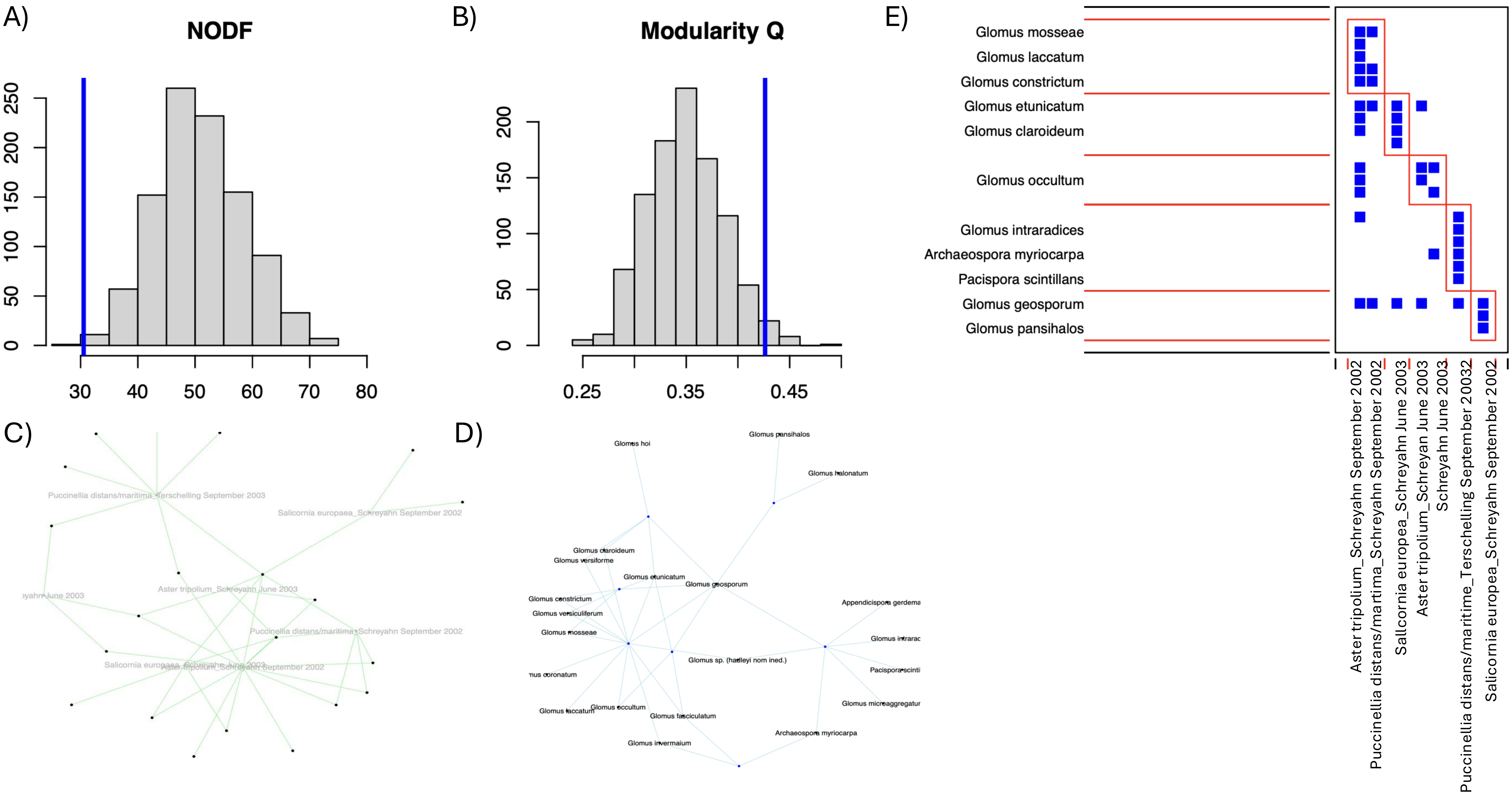

Zhao, Z., Li, X., Liu, M.F., Merckx, V.S., Saunders, R.M. and Zhang, D. 2021. Specificity of assemblage, not fungal partner species, explains mycorrhizal partnerships of mycoheterotrophic Burmannia plants. The ISME Journal 15(6), 1614-1627.

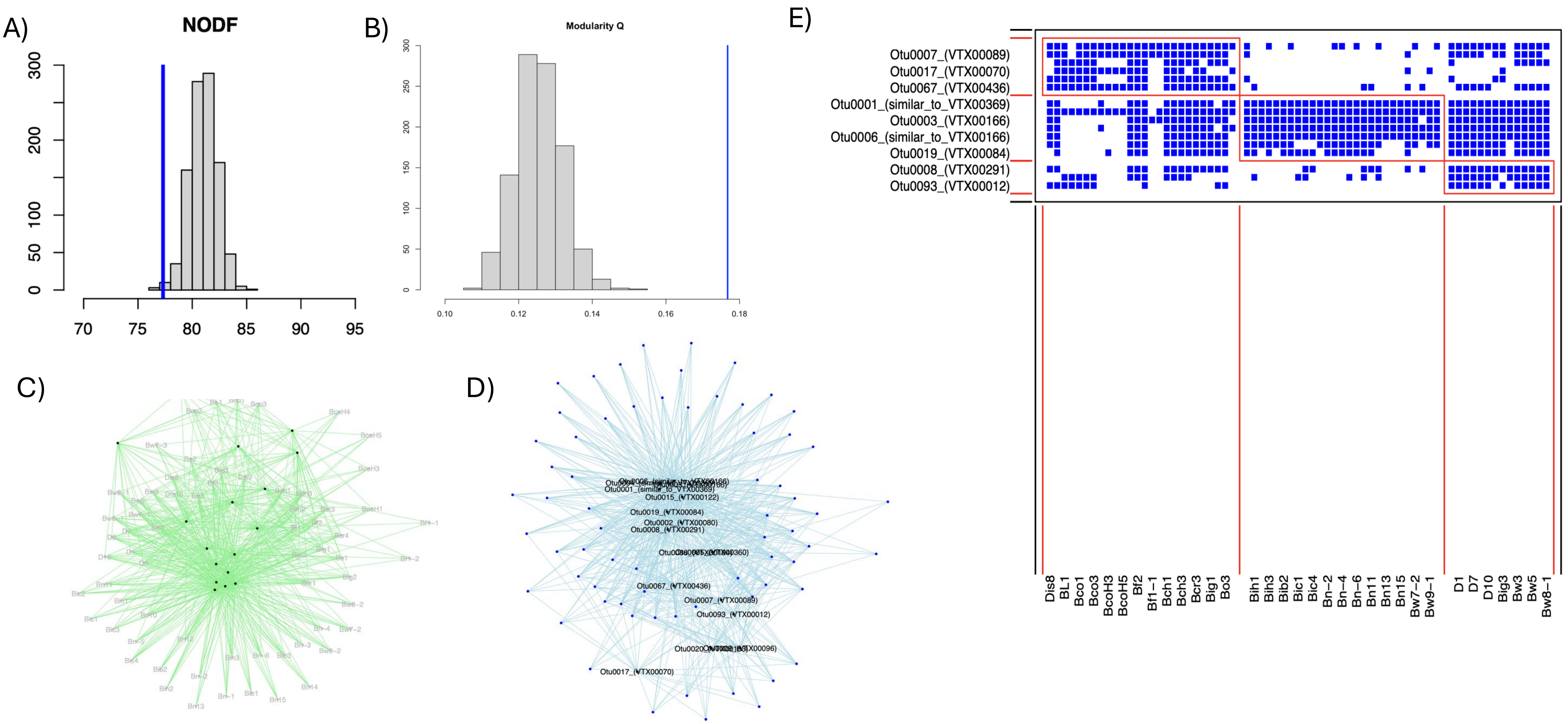
